# Umibato: estimation of time-varying microbial interaction using continuous-time regression hidden Markov model

**DOI:** 10.1101/2021.01.28.428580

**Authors:** Shion Hosoda, Tsukasa Fukunaga, Michiaki Hamada

## Abstract

**Motivation:** Accumulating evidence has highlighted the importance of microbial interaction networks. Methods have been developed for estimating microbial interaction networks, of which the generalized Lotka-Volterra equation (gLVE)-based method can estimate a directed interaction network. The previous gLVE-based method for estimating microbial interaction networks did not consider time-varying interactions.

**Results:** In this study, we developed <u>**u**</u>nsupervised learning based <u>**m**</u>icrobial <u>**i**</u>nteraction inference method using <u>**Ba**</u>yesian es<u>**t**</u>imati<u>**o**</u>n (Umibato), a method for estimating time-varying microbial interactions. The Umibato algorithm comprises Gaussian process regression (GPR) and a new Bayesian probabilistic model, the continuous-time regression hidden Markov model (CTRHMM). Growth rates are estimated by GPR, and interaction networks are estimated by CTRHMM. CTRHMM can estimate time-varying interaction networks using interaction states, which are defined as hidden variables. Umibato outperformed the existing methods on synthetic datasets. In addition, it yielded reasonable estimations in experiments on a mouse gut microbiota dataset, thus providing novel insights into the relationship between consumed diets and the gut microbiota.

**Availability:** The C++ and python source codes of the Umibato software are available at https://github.com/shion-h/Umibato

## 1 Introduction

Comprehensive investigations of microbial community structure using metagenomic analysis have shown that the microbiota plays a key role in humans (Turnbaugh *et al.*, 2007; Wang and Jia, 2016) and the natural environment (Sunagawa *et al.*, 2015; Fierer, 2017; Sunagawa *et al.*, 2020). Microbial interactions, one of the elements of the community structure, are considered as the main drivers of metabolic dynamics (Embree *et al.*, 2015) and have been suggested to influence the host’s health (Fraune *et al.*, 2015). Therefore, microbial interactions have received attention as an important research subject (Phelan *et al.*, 2012; Li *et al.*, 2016). Two main methods are employed for estimating microbial interactions using metagenomic data: correlation-based methods and generalized Lotka-Volterra equation (gLVE)-based methods.

Correlation-based methods evaluate the co-variation of the abundance of each microbe and estimate a positive or negative relationship between the two microbes. Numerous correlation-based methods have been proposed (Faust *et al.*, 2012; Friedman and Alm, 2012; Kurtz *et al.*, 2015; Fang *et al.*, 2015; Ban *et al.*, 2015; Biswas *et al.*, 2016). Recently, McGregor *et al.* (2020) developed the MDiNE, a probabilistic model based on partial correlation coefficients. They applied MDiNE to a human gut microbiota dataset, which included information of patients with Crohn’s disease, and observed differences between cases and controls in interaction networks. An advantage of correlation-based methods is that they can be used for cross-sectional data and have a wide application range. However, the information obtained by this method is limited because it cannot be used to estimate the direction of microbial interactions. The direction of interaction indicates the relationship between microbes, such as symbiosis and competition, and allows a detailed understanding of microbial dynamics (Hibbing *et al.*, 2010; Seipke *et al.*, 2012; Li *et al.*, 2016; Attar, 2016).

The gLVE-based method estimates directed microbial interaction networks by evaluating the contribution of one to the growth of the other. The gLVE-based method represents the microbial growth rate as a linear combination of the abundance of microbes in the community. The growth rates have to be statistically estimated; this estimation requires time-series microbiome data with quantitative abundance obtained by experiments such as qPCR. Nevertheless, the gLVE-based method has been widely employed to elucidate the dynamics using directional information (Stein *et al.*, 2013; Fisher and Mehta, 2014; Buffie *et al.*, 2015; Coyte *et al.*, 2015; Gao *et al.*, 2018; Gibson and Gerber, 2018; Li *et al.*, 2019). Bucci *et al.* (2016) developed MDSINE, a method for estimating directed microbial interaction networks considering perturbations based on gLVE. They estimated the bacteria that inhibit the growth of *Clostridium difficile* and suggested new strategies for the rational design of probiotic cocktails.

A critical limitation of conventional gLVE-based methods is their inability to account for temporal changes in the microbial interaction networks. Microbial interactions have been reported to differ in environments that contain different nutrients (Embree *et al.*, 2015). In environments where nutrition is expected to change dramatically, such as in the human gut, where the nutrition is influenced by diet (Kolodziejczyk *et al.*, 2019), microbial interactions are expected to be time-varying. However, conventional gLVE-based methods cannot estimate such time-varying microbial interactions because they implicitly assume that the microbial interaction network is same at all times.

In this study, we developed <u>**u**</u>nsupervised learning based <u>**m**</u>icrobial <u>**i**</u>nteraction inference method using <u>**Ba**</u>yesian es<u>**t**</u>imati<u>**o**</u>n (Umibato) for estimating time-varying directed microbial interaction networks. We proposed a novel Bayesian model called the continuous-time regression hidden Markov model (CTRHMM), which was included in Umibato. The Umibato algorithm comprises the following two steps:

1. Gaussian process regression (GPR) estimates the growth rates from time-series quantitative microbial abundances (Section 2.3).
2. CTRHMM estimates the time-varying microbial interaction networks from estimated growth rates and time-series quantitative microbial abundances (Section 2.4).

The growth rates estimated by GPR are passed to the CTRHMM together with the estimation uncertainty. This procedure allows for the estimation of directed microbial interactions, considering the uncertainty of growth rate estimation. In addition, CTRHMM has the advantage of being reasonably applicable to data with irregular sampling intervals because it assumes a continuous-time Markov chain. We first confirmed the effectiveness of the Umibato algorithm in synthetic datasets (Section 3.1). We then applied the Umibato algorithm to the mouse gut microbiome dataset and observed that Umibato estimated the specific interaction network in a low-fiber diet (Section 3.2). These results suggest that Umibato can capture changes in the microbial interaction network.

## 2 Materials and methods

### 2.1 Generalized Lotka-Volterra equation

The gLVE, a differential equation describing the symbiosis and competition relationship of microbes, is formulated as follows:

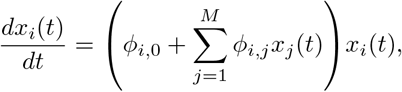

where *x_i_*(*t*) is the quantitative abundance of the *i*-th microbe at time *t*, *ϕ_i,j_*(*j* > 0) is the interaction parameter from the *j*-th microbe to the *i*-th microbe, *ϕ*_*i*,0_ is the growth parameter of the *i*-th microbe, and *M* is the number of different microbes. We defined gLVE parameters as interaction and growth parameters. For computational convenience, gLVE is transformed into the following form:

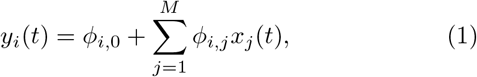

where

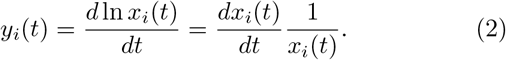

We call *y_i_*(*t*) the growth rates in this paper.

### 2.2 Overview of Umibato algorithm

The Umibato algorithm introduces time-varying gLVE parameters *ϕ_i,j_*(*t*) into Eq. (1): To represent several different conditions caused by environmental events (*e.g.*, nutrient depletion and compound surges due to host diet), we assume that microbial interactions change discretely. Therefore, we defined interaction states as categorical variables that determine the discrete state of interaction. We used 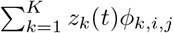 as *ϕ_i,j_*(*t*), where *z_k_*(*t*) is a stochastic process that has a value of 1 when the interaction state at time *t* is the *k*-th state and 0 otherwise; *ϕ_k,i,j_*(*j* = 0) and *ϕ_k,i,j_*(*j* > 0) are the growth and interaction parameters when the interaction state is the *k*-th state, respectively; and *K* is the number of interaction states. Then, the Umibato probabilistic model is based on the following equation:

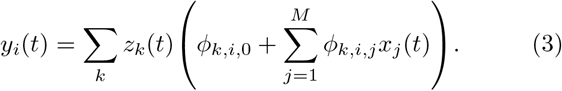

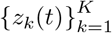 are modeled using a continuous-time Markov chain. The detailed generative processes are described in Section 2.4.1. To use Eq. (3) as a statistical model, we replaced *x_i_*(*t*), *y_i_*(*t*), and *z_k_*(*t*) with quantitative abundance data 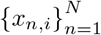, growth rate data 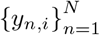, and latent state binary variables 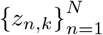, respectively, where *N* is the number of observations. We then obtained the following statistical model:

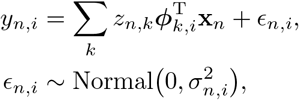

where **x**_*n*_ = (1, *x*_*n*,1_, …, *x_n,j_*, …, *x_n,M_*)^T^ is the quantitative abundance vector of microbes at the *n*-th observation point; ***ϕ**_k,i_* = (*ϕ*_*k,i*,0_, …, *ϕ_k,i,j_*, …, *ϕ_k,i,M_*)^T^ is the gLVE parameter for the *i*-th microbe in the *k*-th state; *ε_n,i_* is an error term of *y_n,i_*; 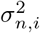 is the variance of *ε_n,i_*; and Normal (*μ, σ*^2^) denotes the normal distribution with a mean of *μ* and a variance of *σ*^2^.

Umibato estimates the time-varying gLVE parameters through the following two steps. First, Umibato estimates *y_n,i_* and 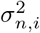 from {*x_n,i_*}*_n_* for the *i*-th microbe using a Gaussian process regression (*cf.* Section 2.3). Second, Umibato estimates {*z_n,k_*}*_n,k_* and {*ϕ_k,i,j_*}*_k,i,j_* from {*x_n,i_*}*_n,i_*, {*y_n,i_*}*_n,i_*, and 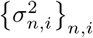 using a continuous-time regression hidden Markov model (*cf.* Section 2.4). We describe the detailed Umibato procedure in Algorithm 1 (the notations are described in Supplementary Table S1).

### 2.3 Gaussian process regression for growth rate estimation from time-series quantitative microbiome data

GPR is a probabilistic regression model based on a kernel function. We used GPR to estimate the growth rates and their variances from quantitative abundances. An essential property of GPR is its ability to estimate the distribution of the true function. In Umibato, the logarithmic abundance trajectories of microbes are estimated as true functions, and *y_n,i_* and 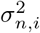 are given as follows:

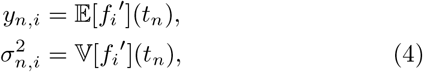

where *f_i_* is the logarithmic abundance trajectory function of the *i*-th microbe; *f_i_*′ is the growth rate function, that is, the time derivative of the estimated *f_i_* (Eq. (2)); 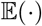 and 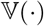 are the expectation and variance with respect to the posterior distribution of *f_i_*, respectively; and *t_n_* is the time of the *n*-th observation point. 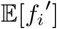 was obtained by differentiating the kernel function, and 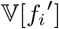 was approximated by sampling *f_i_*′ from the posterior distribution.

Preliminary experiments have shown that the growth rate estimation by GPR is more powerful than that by penalized spline interpolation, which has been used in previous studies (Supplementary Figure S1). In addition, GPR has another advantage in that hyperparameters, such as a penalty coefficient in penalized spline interpolation, are not required. Note that we used the term “hyperparameter” here as a parameter that was not estimated in iterations.

#### 2.3.1 Parameter estimation

We estimated the kernel parameters using the maximum likelihood estimation. We used a radial basis function kernel and Gaussian noise and implemented GPR parameter estimation using the Python library GPy (https://github.com/SheffieldML/GPy). To avoid the logarithm of zero, we replaced zeros in the abundance matrix by a pseudo abundance, where the pseudo abundance is set to be the largest 10^*r*^ that does not exceed the minimum non-zero value of the abundance matrix (*r* is an arbitrary integer).

#### 2.3.2 Outlier detection

In the growth rate estimation, we excluded the observation points that were not in the middle 90% of the distribution estimated by the GPR as outliers. We then conducted GPR again, and the results were used as the final estimates.

#### 2.3.3 Calculating the variance of the estimated growth rate

We sampled *f_i_* from the posterior distribution estimated by GPR 100 times and calculated the unbiased variance. Variances below 10^−4^ were corrected to 10^−4^ because very small variances would interfere with the estimation of the CTRHMM.

### 2.4 Continuous-time regression hidden Markov model for estimating time-varying gLVE parameters

We proposed a novel Bayesian probabilistic model, CTRHMM. CTRHMM was used to estimate the interaction states and the corresponding networks using growth rates and their variances estimated by GPR. CTRHMM is a model similar to the input-output hidden Markov model (Bengio and Frasconi, 1994) but differs in that a continuous-time Markov chain assumes the states. The continuous-time Markov chain enables the application of the model to data with irregular sampling intervals. This ability is useful because periodic sampling of the microbiome is difficult in some cases. For example, the sampling interval for human gut microbiota studies depends on the defecation interval. The continuous-time Markov chain, which can consider sampling intervals, is therefore more suitable for modeling interaction states than the discrete-time Markov chain. In Section 2.4.2, we introduce a variational inference procedure for CTRHMM. To the best of our knowledge, this is the first time variational inference has been applied to continuous-time hidden Markov models.

#### 2.4.1 Generative process of CTRHMM

The generative process of CTRHMM is given as follows:

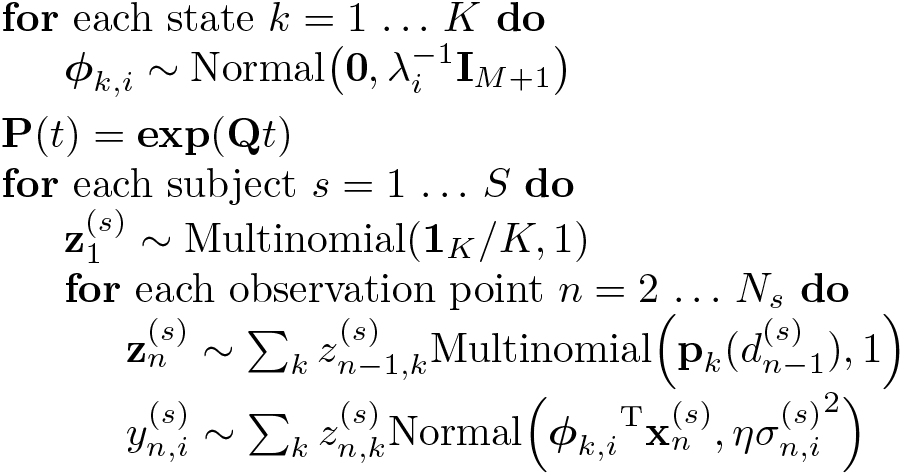

Here, *λ_i_* is the precision parameter for the prior distribution of gLVE parameters to the *i*-th microbe; **I**_*M*+1_ is the (*M* + 1)-dimensional identity matrix; **P**(*t*) = **p**_1_(*t*), …, **p**_*k*_(*t*),…, **p**_*K*_(*t*))^T^ is the transition probability matrix for the elapsed time *t*; **exp**(·) is the matrix exponential function; **Q** is the transition rate matrix; *S* is the number of subjects; .^(*s*)^ denotes the *s*-th subject; 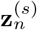 is a one-hot vector indicating the interaction state of the *n*-th observation point of the *s*-th subject; Multinomial(**p**, *N*) denotes the multinomial distribution with the event probability vector **p** and number of trials *N* ; **1**_*K*_ is a *K*-dimensional vector in which all the elments are 1; *N_s_* is the number of observation points of the *s*-th subject; 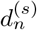 is the time interval of the *s*-th subject between the *n*-th observation point and the (*n* + 1)-th observation point; and *η* is the reciprocal of the *inverse temperature* in the field of statistical physics, which adjusts for the inuence of likelihood and prior distribution. Figure 1 shows a graphical representation of CTRHMM.

**Figure 1:**
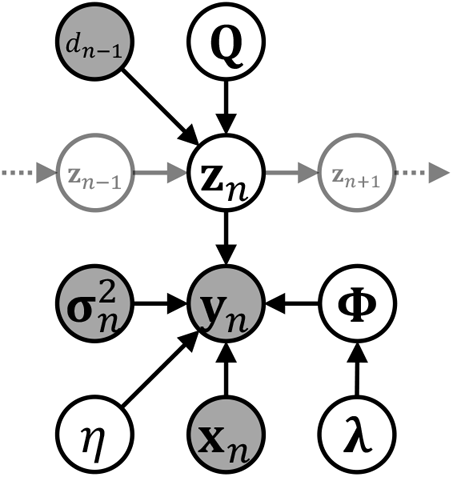
Graphical representation of the proposed model (CTRHMM). Only the *n*-th observation point is displayed, and the subject indices are omitted. The shaded and white circles represent the observed and unobserved variables respectively.

#### 2.4.2 Variational inference for estimating CTRHMM parameters

We present a variational inference procedure for CTRHMM. Variational inference is a parameter estimation method for Bayesian probabilistic models that introduces an approximate posterior distribution and maximizes the evidence lower bound (ELBO) (Attias, 2000). Maximizing ELBO is equivalent to minimizing the Kullback-Leibler divergence between an approximate posterior distribution and the true posterior distribution. We introduce the following approximate posterior distributions:

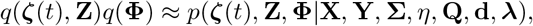

where **X**, **Y**, **Σ**, and **Z**, are the quantitative abundance matrix, growth rate matrix, growth rate variance matrix, and onehot state matrix of all subjects at all timepoints, respectively; ***ζ***(*t*) is the stochastic process of states between observation points; **Φ** is the gLVE parameter tensor; **d** is the time interval vector between each observation point of all subjects; ***λ*** is the parameter vector for the prior distribution of **Φ**; and *N_s_* is the number of observation points of the *s*-th subject. In addition, we assume that the following factorization is possible for the approximate posterior distribution:

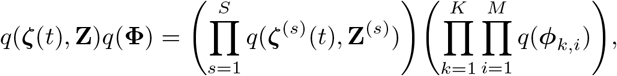

where ***ϕ**_k,i_* is the gLVE parameter for the *i*-th microbe in interaction state *k*. The detailed notations are described in Supplementary Table S1. Then, ELBO 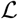 is written as

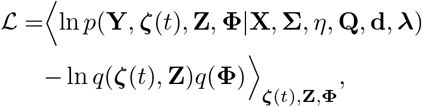

where 〈·〉_**ζ**(*t*),**Z**,**Φ**_ denotes the expectation with respect to *q*(***ζ***(*t*), **Z**)*q*(**Φ**). We maximize 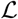 in the variational inference. Using the variational method, *q*(***ϕ***_*k,i*_) satisfying 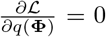 is obtained as follows:

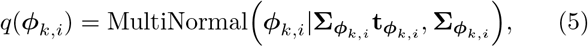

where MultiNormal(***μ***, **Σ**) denotes the multivariate normal distribution with mean vector ***μ*** and covariance matrix **Σ**, and

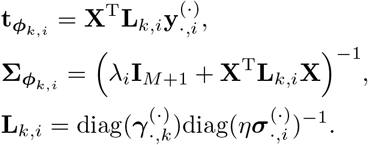

Here, diag(**a**) is the diagonal matrix whose diagonal elements are **a**, 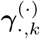 is the expected vector of the *k*-th column of **Z** with respect to *q*(**Z**), 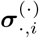 is the *i*-th columns of **Σ**, and 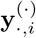 is the *i*-th columns of **Y**. Similarly, *q*(**Z**^(*s*)^) is obtained as follows:

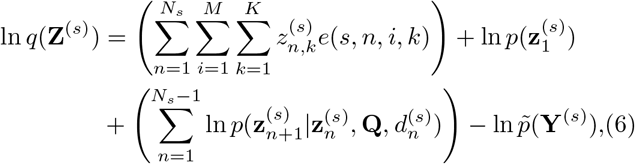

where 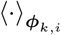 denotes the expectation with respect to *q*(***ϕ***_*k,i*_), ln 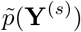 is the normalization constant for *q*(**Z**^(*s*)^), and

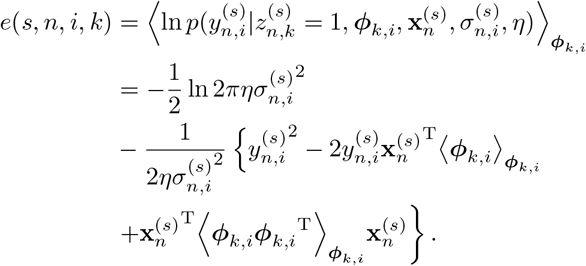

The second and third terms on the right-hand side of Eq. (6) are given by

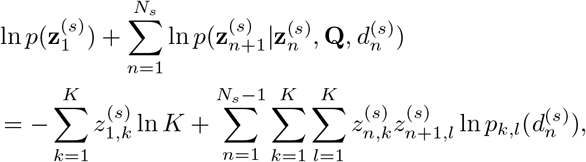

where *p_k,l_*(*t*) is the transition probability from the *k*-th state to the *l*-th state for the elapsed time *t*. Eq. (6) can be computed using the forward–backward algorithm (Baum *et al.*, 1972). We used the true conditional posterior distribution 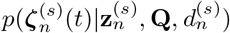 as 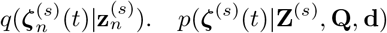 can be expressed as follows:

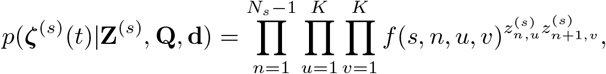

where

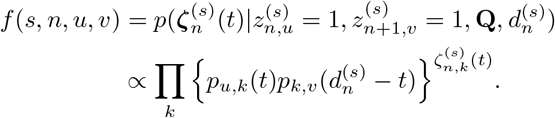

The implementation does not calculate the distribution but only the following expectations:

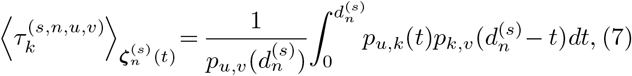

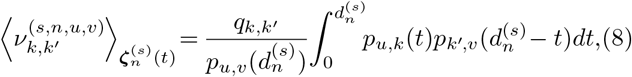

where 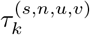 and 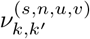 are the time interval of state *k* and the number of transitions from state *k* to *k*′ between the *n*-th and (*n* + 1)-th observation points of the *s*-th subject when 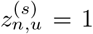 and 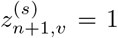 hold, respectively; 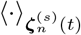 denotes the expectation with respect to 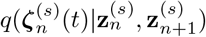; and *q_k,k_*′ is the (*k, k*′) element of the transition rate matrix **Q**. These expectations can be obtained by calculating one matrix exponential according to a previous study on a continuous-time hidden Markov model (Liu *et al.*, 2015). Type II maximum likelihood estimation of the other parameters were obtained as follows:

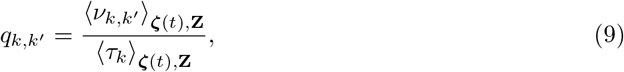

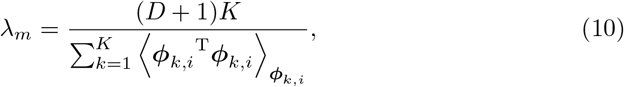

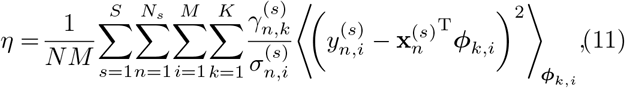

where 〈·〉_**ζ**(*t*),**Z**_ denotes the expectation with respect to *q*(***ζ***(*t*), **Z**); 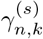 is the expectation of 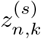 with respect to 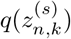 and *τ_k_* and *ν_k,k_′* are the time interval of state *k* and the number of transitions from state *k* to *k*′ at all periods, repectively 〈*τ_k_*〉 _**ζ**(*t*), **Z**_ and 〈 *ν*_*k,k*_′ 〉 _**ζ**(*t*),**Z**_ can be calculated from 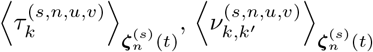, and the expected number of transitions between the observation points obtained by the forward–backward algorithm. Finally, ELBO can be rewritten as

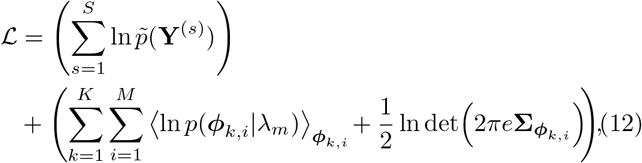

when Eq. (6) holds, where det(·) denotes the matrix determinant calculation. A detailed derivation of the variational inference algorithm is provided in Supplementary Section S1.

#### 2.4.3 State deletion

In the process of estimating the CTRHMM parameters, only a small number of observations are assigned to one state. A state that is allocated only the number of observation points less than or equal to *M* + 1 is overfitting because it can reconstruct the observation points without any errors. We introduced heuristics to remove such states and the corresponding parameters during estimation. In detail, we deleted a state that satisfies the following conditions:

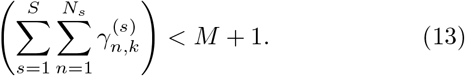

The heuristics were performed so that at most one cluster was deleted per iteration. In the next iteration, where this operation is performed, convergence is not determined because ELBO may decrease.

#### 2.4.4 Convergence determination

The learning process is terminated when the change in the ELBO between the previous and current steps is less than 10^−4^, or when the total number of iterations exceeds 100.

#### 2.4.5 Standardization of quantitative abundance matrix

The quantitative abundance matrix **X** was standardized such that the mean is 0 and the variance is 1 for each microbe to equalize the effect of the prior distribution of **Φ**. Standardization of **X** can also remove the 16S rRNA gene copy number bias when comparing the interaction parameters to a microbe. In other words, standardization discards information regarding the scale of microbial abundances and enables the estimation of contributions to growth rates that can be compared (*cf.* Figure 5). In a synthetic data experiment (Section 3.1), we corrected the gLVE parameters estimated using standardized **X** to compare them with the true parameters. The detailed procedure is described in Supplementary Section S2.

### 2.5 Synthetic data experiment

We generated gLVE parameters using a generative process and prepared three different synthetic datasets according to gLVE. The generative process is the same for the three datasets and was set up based on the MDSINE assumptions regarding signs (MDSINE assumes *ϕ_i,j_* < 0(*i* = *j*) and *ϕ*_*i*,0_ > 0). Each dataset has a different combination of the number of states *K* (1 or 2) and the number of microbes *M* (5 or 10). The details of the settings of states and parameters are described in Supplementary Section S3. We conducted 100 CTRHMM trials for each initial number of states *K_init_* = 1, …, 5. We adopted a trial with the highest ELBO maximized in the learning process. The performances of MDSINE and Umibato were evaluated using the two measurements: the Pearson’s correlation coefficients and the mean absolute error (MAE). Pearson’s correlation coefficients and MAE were calculated between the true and estimated parameters at each observation point. To evaluate Umibato, we compared the true parameter with the parameter corresponding to the state of the maximum likelihood path for each observation point.

#### Algorithm 1

The Umibato algorithm

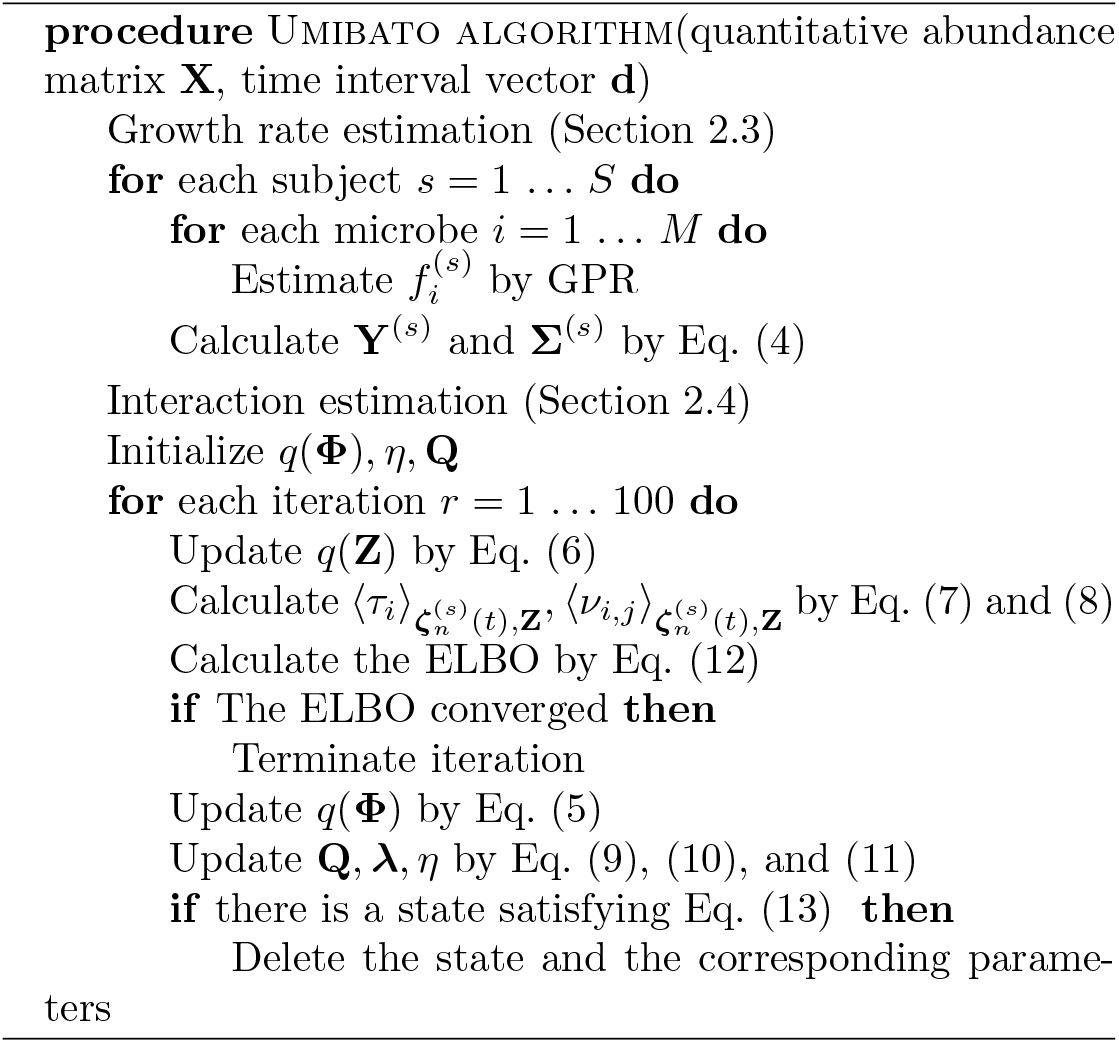

### 2.6 Real data experiment

We used the time-series quantitative mouse gut bacterial dataset of Bucci *et al.* (2016), which was obtained from the feces of seven mice that had orally ingested 13 strains of *Clostridium*. These 13 bacterial strains were determined based on the work of Atarashi *et al.* (2013), which suggested that the strains induced T_reg_ cells. Supplementary Table S2 shows the lineage of each strain. The data were measured using strain-specific qPCR primers. All mice were fed a high-fiber diet, five of the seven mice were temporarily switched to a low-fiber diet, and two of the seven mice were not switched and acted as controls. The first five mice had 56 observation points each, and the latter two mice had 25 observation points each. We downloaded this dataset from the MDSINE software repository (https://bitbucket.org/MDSINE/mdsine/downloads). We conducted 10000 CTRHMM trials for each *K_init_* = 1, …, 15. We adopted a trial with the highest ELBO maximized in the learning process.

## 3 Results

### 3.1 Accuracy evaluation on synthetic dataset

We tested the performance of Umibato compared with that of MDSINE on synthetic datasets generated using known parameters according to gLVE (*cf.* Section 2.5). Four algorithms, BAL, BVS, MLRR, and MLCRR, were implemented in MDSINE (Bucci *et al.*, 2016), and the performance of Umibato was compared with those of all others. The performance of Umibato was visualized over two settings: (1) the case in which the number of true states was given as *K_init_* (called “true model case”) and (2) the case in which several *K_init_* were used similar to the real data experiment (called “practical case”).

Figure 2 shows the means of the correlation coefficients between true parameters and parameters estimated by each method for each synthetic dataset. First, Dataset 1 is a single-state dataset; that is, it obeys the equations assumed in previous studies. The mean correlation coefficients of Umibato in the true model case and practical case and those of BAL, BVS, MLRR, and MLCRR in Dataset 1 were 1.0, 0.90, 0.32, 0.45, 1.0, and 1.0, respectively. Umibato exhibited high performance in both cases. The estimation performance of Umibato in the practical case was lower than that in the true model case because the state estimation of some of the observation points failed (Supplementary Figure S2). MLRR and MLCRR showed high performance in Dataset 1 because Dataset 1 was a single state. Second, Dataset 2 was generated by two states. The mean correlation coefficients of Umibato in the true model and practical cases and those of BAL, BVS, MLRR, and MLCRR in Dataset 2 were 0.99, 0.90, 0.16, 0.33, 0.53, and 0.56, respectively. The performance of Umibato on Dataset 1 was similar to that on Dataset 1; however, MLRR and MLCRR were much less accurate for Dataset 1 than for Dataset 2. Previous gLVE-based methods were suggested not to accurately estimate microbial interaction networks for multi-state data such as Dataset2. For state estimation, Umibato in the true model case failed for only four observation points, whereas Umibato in the practical case failed for 88 observation points (Supplementary Figure S3ac). Third, In Dataset 3, the number of different microbes was doubled compared to those in Datasets 1 and 2. The mean correlation coefficients of Umibato in the true model and practical cases and those of BAL, BVS, MLRR, and MLCRR in Dataset 3 were 0.80, 0.89, 0.17, 0.14, 0.19, and 0.25, respectively. The estimation performance of Umibato in the true model case was lower for Dataset 3 than that for Dataset 2, and in the practical case, Umibato performed better than in the true model case. For state estimation, Umibato in the true model case did not fail for any of the observation points, whereas Umibato in the practical case failed for 22 observation points (Supplementary Figure S3bd). In summary, MDSINE showed high performance on Dataset 1 but low performance on Datasets 2 and 3 because MDSINE assumed a single interaction in the microbiota, whereas Umibato showed high performance on all datasets. The tendencies discussed in this subsection can be observed in the MAE evaluation (Supplementary Figure S4).

**Figure 2:**
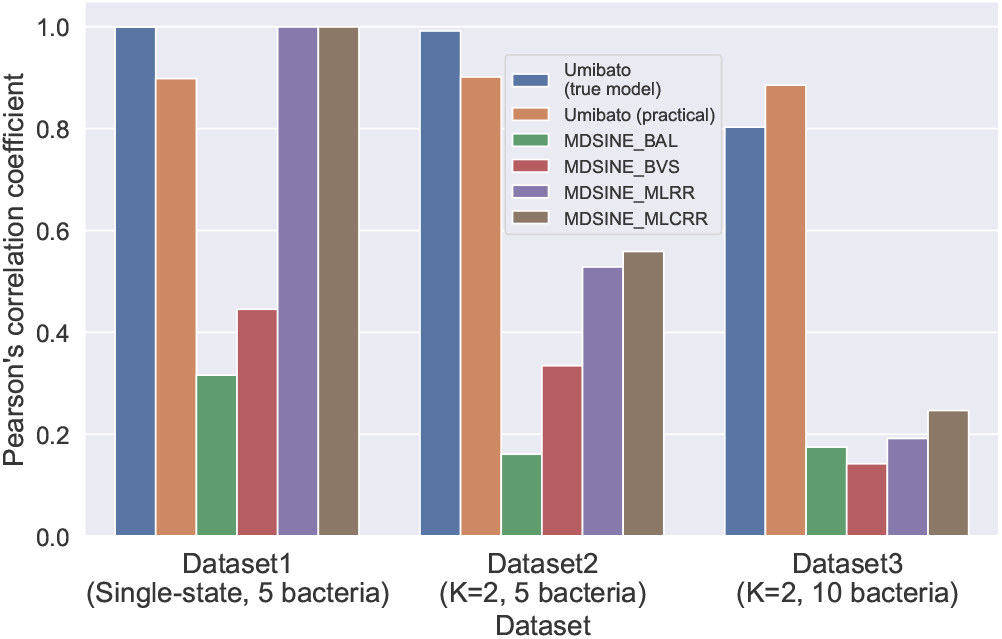
Pearson’s correlation coefficients between the true parameters and parameters estimated by each method for each synthetic dataset. The *x*- and *y*-axes indicate the mean of the Pearson’s correlation coefficients of gLVE parameters of all observation points for the datasets. The six bars indicate Umibato in the true model case, Umibato in the practical case, BAL, BVS, MLRR, and MLCRR in order from left to right. The “true model case” and “practical case” are described in Section 3.1.

### 3.2 Results on real mouse gut microbiome dataset

We applied Umibato to the mouse gut bacterial dataset and found that the maximized ELBO for a trial with *K_init_* = 6 was the highest across all trials (Supplementary Figure S5). The final number of states was 5, and we will discuss the results of this trial in further analysis. As the estimated interaction parameters, we used the expectation of **Φ** with respect to *q*(**Φ**). The computational time was described in Supplementary Section S4.

#### 3.2.1 Estimated interaction state trajectories

To see how the bacterial interaction states changed, we examined the maximum likelihood paths on the mouse gut microbiota dataset. Figure 3 shows the estimated state paths and dietary information of the mice. We found that the State 5 was frequently estimated on low-fiber diet days and not in the control mice. These results suggest that the switch from the high-fiber to low-fiber diets altered bacterial interactions in the mouse gut. States 1 and 2 were frequently estimated just after the first day of the experiment and may be unstable interactions caused by orally ingested bacteria. The transitions between States 3 and 4 were frequently observed. This result suggested that the states were not constant even with the same diet (*i.e.*, the high-fiber diet). To verify the robustness of these observations, we also visualized the results for *K_init_* = 3. We chose *K_init_* = 3 because States 3, 4, and 5 were mainly estimated. The same tendency was observed for the results estimated with *K_init_* = 3 (Supplementary Figure S6).

**Figure 3:**
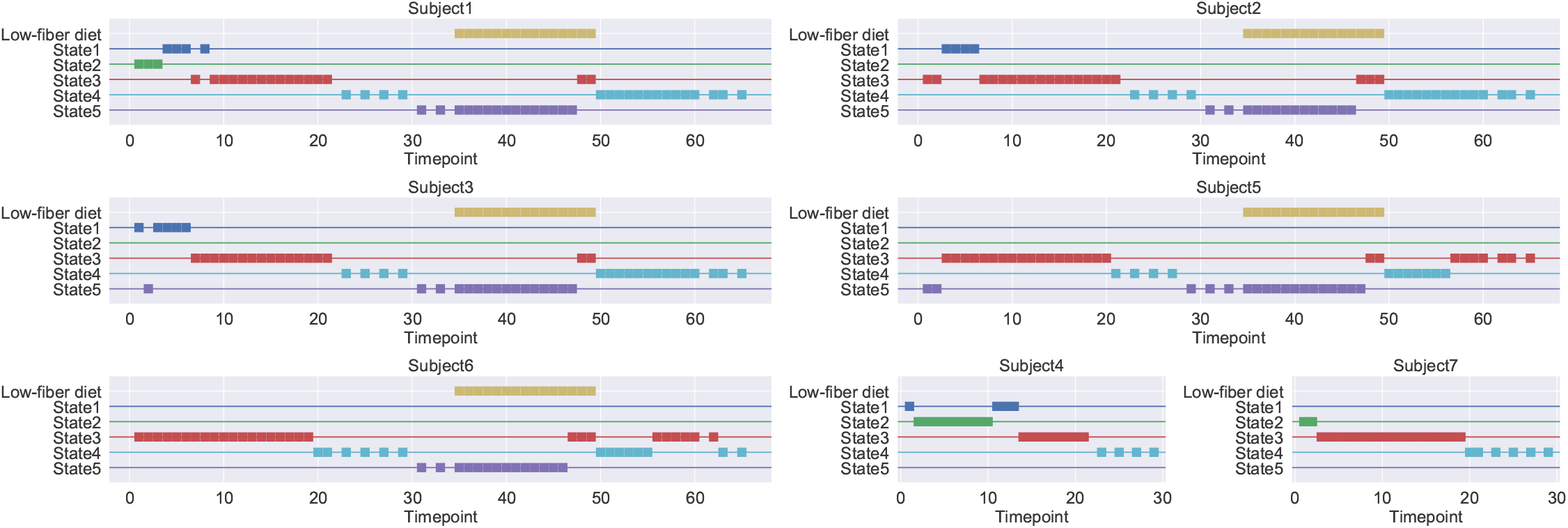
Interaction state path estimated by Umibato on the mouse gut bacterial dataset. Each axis indicates each mouse gut microbiome. The *x*- and *y*-axes indicate the days since the start of the experiment and the interaction states and low-fiber diet days, respectively. Each square marker indicates the state estimated at the time point. For example, there is a marker in (10, State3) in the figure of Subject 1, and it indicates that the interaction state of day 10 of subject1 is State 3. Subjects 4 and 7 were controls; that is, they were fed a high-fiber diet at all time points.

#### 3.2.2 Estimated transition rate matrix of interaction state

To investigate the relationship between the states, we next examined the transition rate matrix **Q**. Figure 4a shows the value of each element of **P**(1), that is, the transition probability matrix of the state after one day. All diagonal components are above 0.5, indicating that each state is likely to last for more than one day. State 1 has a high probability of transition to State 3. States 1 and 3 may be the intermediate states between the states after oral ingestion of bacteria and the stable states. Figure 4b shows the value of each element of **P**(7), that is, the transition probability matrix of the state after one week. The diagonal components of States 3, 4, and 5 were approximately 0.5, whereas the diagonal components of States 1 and 2 were 0.052 and 0.12, respectively. These results show that States 1 and 2 are short-term states that are likely not to last for a week. States 3 and 4 have a high probability of transition to the other state, which suggests that these states represent interaction networks stabilized by mutual transitions. Note that the probability of each diagonal component includes the probability of transitioning back in time.

**Figure 4:**
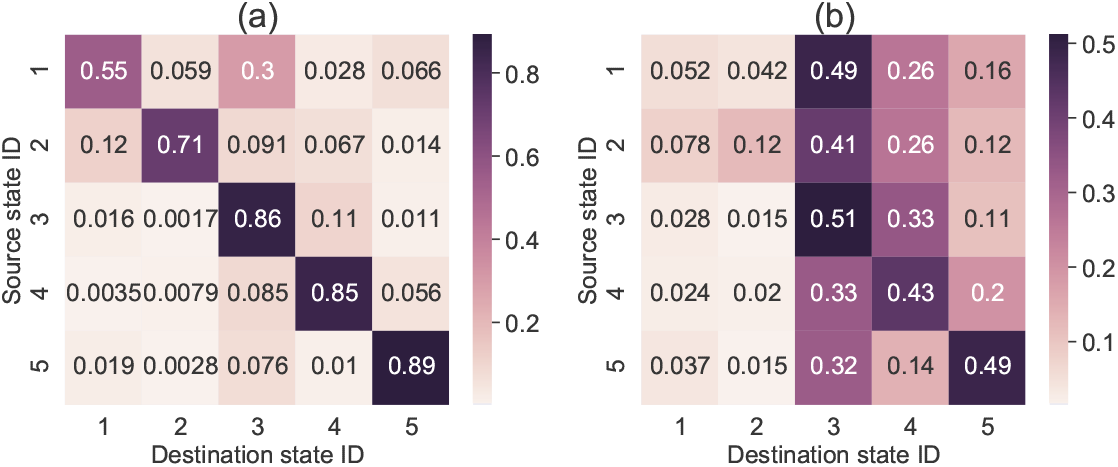
Transition probability matrix **P**(·) of interaction states estimated by Umibato on the mouse gut bacterial dataset. The *x*- and *y*-axes represent the destination and source state IDs, respectively. Darker colors indicate higher probabilities. **(a) P**(1), that is, the transition probability matrix after one day. **(b) P**(7), that is, the transition probability matrix after one week.

#### 3.2.3 Directed interaction networks for each interaction state

To understand the differences in the relationships between the bacteria, we verified the interaction parameters of each state. Figure 5 shows the directed networks based on the estimated interaction parameters divided by the standard deviation of **Y** for the parameters those above the threshold (0.25). We can see several parasitism relationships in State 5. Here, the word “parasitism” refers to a relationship between A and B such that A contributes to the increase in B and B contributes to the decrease in A. In particular, strains 6, 15, and 28 showed three-way parasitism. The interactions of State 4 are active, whereas those of State 3 are relatively inactive. Together with the discussion in Section 3.2.1 and Section 3.2.2, our results suggest that the interaction network frequently switches between dense and sparse states. These 13 strains were suggested to synergistically amplify the induction of T_reg_ cells in a microbial community-dependent manner (Atarashi *et al.*, 2013). Therefore, the processes required for T_reg_ induction may correspond to some or all states. A positive edge from strains 13 to 15 is common to States 3, 4, and 5. This positive edge was also observed in the network estimation of a single-state experiment (Supplementary Figure S7).

**Figure 5:**
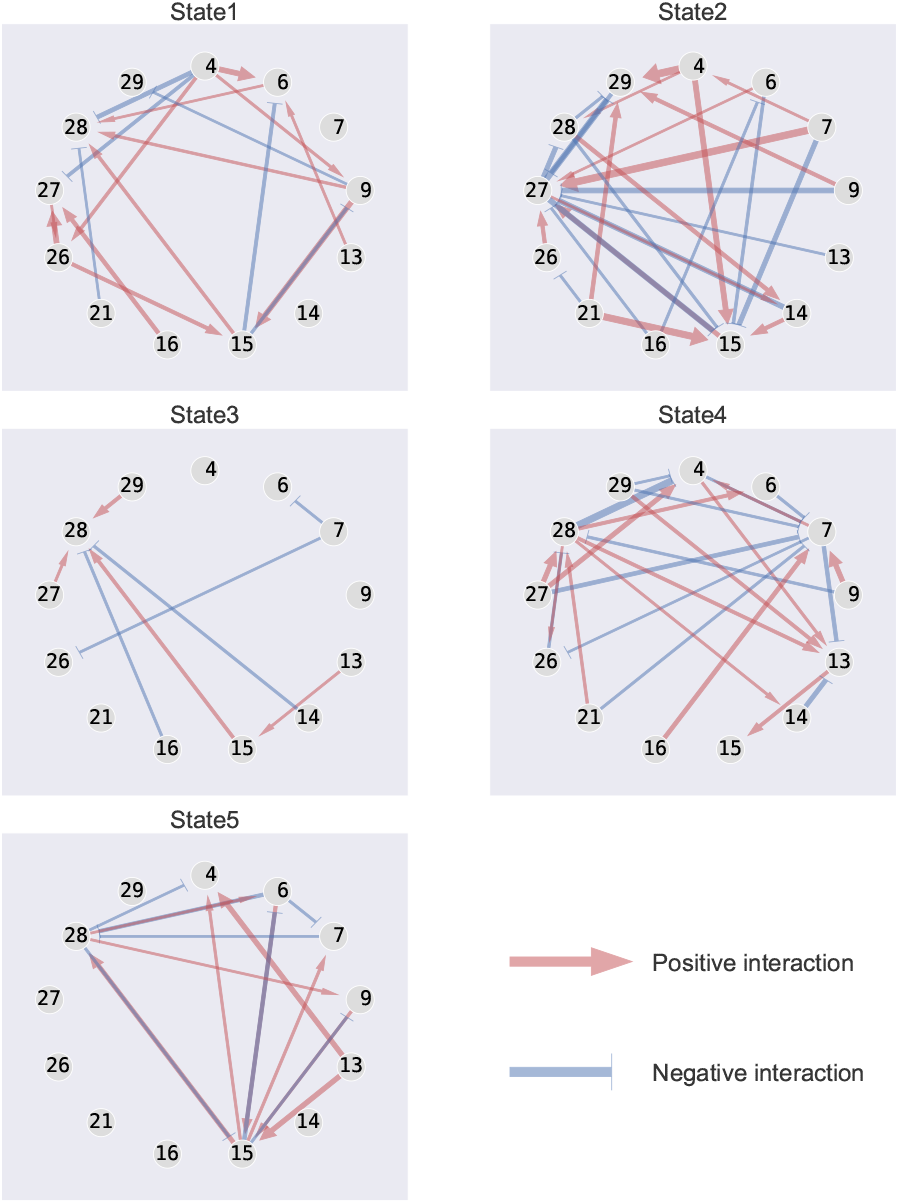
Estimated bacterial interaction networks corresponding to states. The number shown in each node indicates the strain ID in Supplementary Table S2. The width of the edge indicates that the estimated interaction parameter divided by the standard deviation of the growth rate, and values below 0.25 were omitted. The red arrow and blue T-shaped edges indicate positive and negative interactions, respectively.

#### 3.2.4 Simulated bacterial abundance trajectories using estimated parameters

The gLVE enables the simulation of bacterial trajectories. Similarly, the continuous-time Markov chain can simulate state trajectories because of its ability to generate states. To assess the effect of a long-term low-fiber diet on the gut microbiota, we simulated the bacterial abundance trajectory using the estimated parameters. After 20 days in State 5, which is considered the bacterial interaction state on the low-fiber diet, we randomly shifted the state according to estimated **Q**. The procedure for generating bacterial trajectories is the same as that for generating synthetic datasets (Section 2.5) described in Supplementary Section S3, where Gamma(1, 2) was used to generate the initial value of abundance. Figure 6 shows the simulated interaction state and the bacterial abundance trajectories. Strains 13, 26, 27, and 28 increased in State 5 and decreased in States 3 and 4, while strain 21 decreased and increased. Strains 7 and 14 showed an increasing trend in all periods. Slight variation was observed in the abundance of strain 16. Strain 16 may be less affected by the abundance of other bacteria. The abundance of strains 6 and 9 almost disappeared during State 5 and could not be restored even after a long period in States 3 and 4. This result suggests that a long-term low-fiber diet may lead to an irreversible decrease in the diversity of microbiota.

**Figure 6:**
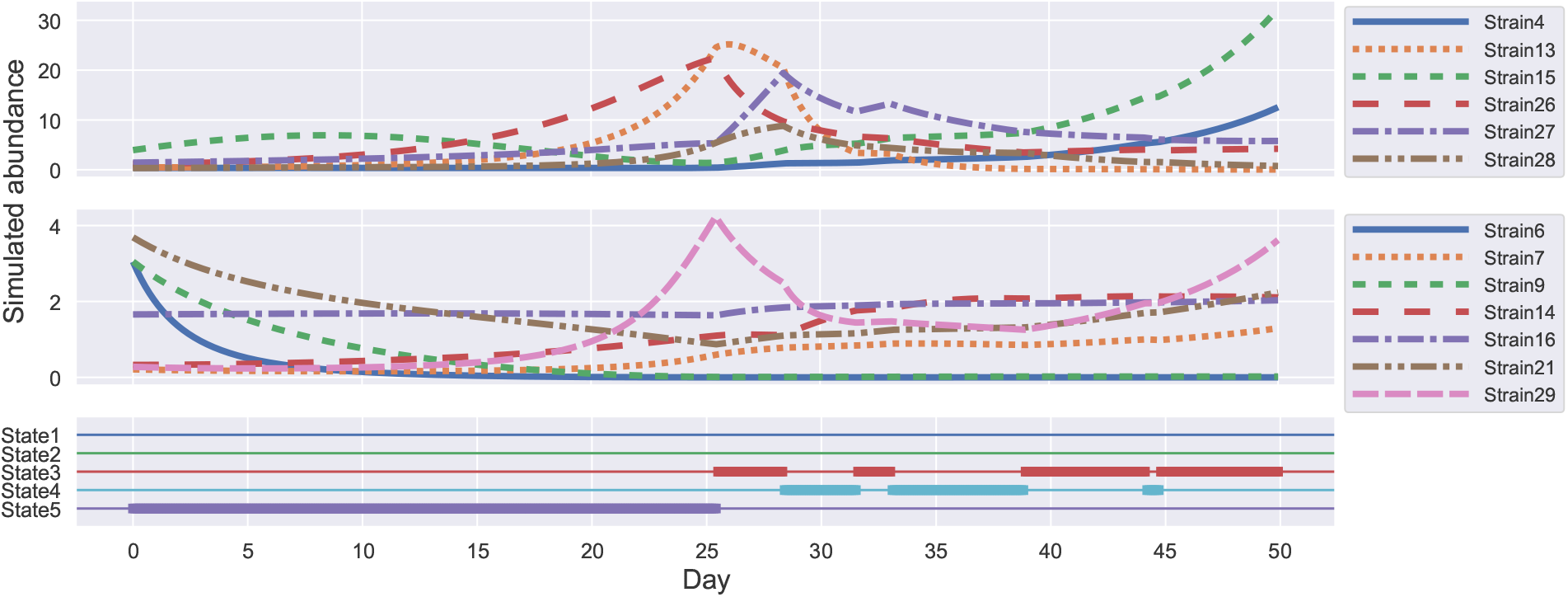
Simulated interaction state and bacterial abundance trajectories. We fixed the first 20 days in State 5 and randomly shifted the state for the next 30 days. The trajectories were divided into two pains because of differences in the abundance scale. The top, middle, and bottom figures show the simulated trajectory of the bacteria with high abundance and low abundance and the simulated interaction states, respectively. The *x*-axis indicates the number of days elapsed, whereas the *y*-axis indicates the abundance of bacteria or the interaction state.

## 4 Discussion

Here, we propose a new method, Umibato, for estimating time-varying microbial interaction networks. The first step of Umibato is growth rate estimation. Our proposed method uses GPR, which enables accurate and hyperparameter-free growth rate estimation. The second step is interaction estimation. Umibato adopted a new probabilistic model, the CTRHMM, proposed in this study. CTRHMM can capture changes of gLVE parameters in time by assuming discrete state variables. Umibato was shown to outperform existing methods for synthetic datasets. In the real mouse gut dataset, specific states were estimated at the low-fiber dietary time, suggesting that gut bacterial interactions changed in a diet-dependent manner.

There is still room for improvement in the growth rate estimation. First, we can utilize trajectory noise as well as variances of growth rates. Umibato estimated growth rate distributions from trajectories of bacterial quantitative abundances and used their expectations and variances in the CTRHMM. By extending CTRHMM, we can also utilize the trajectory noise. Specifically, the true quantitative abundance matrix 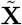 is defined as a new latent variable, and **X** is assumed to be stochastically generated from 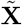. Here, the trajectory noise estimated by the GPR is used as the noise of the distribution of **X**. As suggested by Cao *et al.* (2017), noise exists in the quantitative abundance matrix **X**. Therefore, it may be useful to estimate gLVE parameters considering the trajectory noise. Second, it is also possible to use growth rate estimation methods based on the peak-to-trough ratio (PTR), which has been proposed recently (Korem *et al.*, 2015; Brown *et al.*, 2016; Emiola and Oh, 2018; Suzuki and Yamada, 2020). The PTR-based growth rate estimation methods require a metagenomic shotgun sequencing dataset and are based on the fact that the higher the growth rate, the more DNA is mapped around the replication origin (Cooper and Helmstetter, 1968; Bremer and Churchward, 1977). Adopting the PTR-based growth rate estimation enables using cross-sectional datasets. However, the performance of PTR-based methods is questionable (Long *et al.*, 2020), and the use of this method should be considered carefully.

Three directions can be used to improve the interaction state estimation by CTRHMM. The first direction is model selection. In the synthetic dataset experiments, several observation points (> 20 points out of all 700 points) failed in the state estimation of Umibato in the practical case, whereas Umibato in the true model case could estimate the correct state with few failures (< 5 points out of all 700 points) (Section 3.1). Therefore, estimating the correct number of states in advance, that is, accurate model selection, enables us to improve the state estimation. A typical model selection method involves the use of information criteria. One information criterion that can be applied to mixture models is the WBIC (Watanabe, 2013). However, WBIC assumes independent and identically distributed latent variables and is not applicable to HMMs. A mathematically justified method for applying WBIC to HMMs has not yet been established. The second direction is a Bayesian estimation of the transition rate matrix **Q**. The Bayesian estimation of **Q** is expected to be more robust than the maximum likelihood estimation adopted in this study. Robust estimation of **Q** allows robust state estimation because **Q** constitutes the prior distribution of interaction states **Z**. To perform a Bayesian estimation of **Q** in variational inference, we compute the expectations with respect to the approximation posterior distribution *q*(**Q**) for calculations involving **Q**, such as *q*(**Z**)(Eq. (6)) and *q*(*ζ*(*t*) **Z**)(Eq. (7), (8)), which cannot be computed analytically. Therefore, the approximation of the matrix exponential or sampling approximation of the expectations must be used. The third direction is semi-supervised learning. In the application of mouse gut bacterial dataset (Section 3.2), State 5 interaction, which seems to be due to low-fiber diets, was estimated without giving labels in an unsupervised learning framework. We also suggested that there were two primary states (*i.e.*, States 3 and 4) on high-fiber diets. As we have seen, unsupervised learning is powerful, but we can also take an approach that utilizes labels for learning. In the present case, semi-supervised learning can be applied for state estimation using diet labels. That is, CTRHMM is trained where some of the states of the observation points are known. This method can be easily implemented by fixing 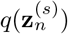 of the observation points given the label during the model learning iterations (Nigam *et al.*, 1998).

There are several candidates for the application of Umibato. First, the most interesting candidate is the human gut microbiota. Relationships between the host’s disease and microbial interactions have been suggested (Fraune *et al.*, 2015; McGregor *et al.*, 2020); hence, changes in microbial interactions estimated by Umibato may reveal the dynamics of contracting diseases. Indeed, simulations of bacterial trajectories (Section 3.2.4) suggested the effect of a long-term low-fiber diet on the gut microbiome. Some bacteria were eliminated by the low-fiber diet. A decrease in community diversity in the human gut microbiome is called dysbiosis and has been reported to be associated with several diseases (Tamboli *et al.*, 2004; Levy *et al.*, 2017). Unfortunately, to the best of our knowledge, there are no long-term quantitative time-series data of human gut microbiota. Therefore, further data accumulation is required. Second, the application of Umibato to the microbiome data of the natural environment, such as ocean and soil, may be useful. Umibato is expected to be effective for analyzing environmental data with dramatic changes in conditions (Ramsby *et al.*, 2018). Investigating the relationship between seasons/weather/temperature and microbial interactions in natural environments through long-term sampling may provide new insights into microbial research.

## Supporting information

Supplementary data including Figures S1, S2, S3, S4, S5, S6, and S7, Tables S1 and S2, and Sections S1, S2, and S3 are available.

## Acknowledgements

The computations in this research were performed using supercomputing facilities at the National Institute of Genetics in the Research Organization of Information and Systems.

## Supplementary data

Supplementary data including Figures S1, S2, S3, S4, S5, S6, and S7, Tables S1 and S2, and Sections S1, S2, S3, and S4 are available.

## Author contributions

SH conceived this study. MH supervised this study. SH designed the algorithms, implemented the method, and performed all the computational experiments. SH, TF, and MH analyzed the results. SH wrote the draft. TF and MH revised the manuscript critically. All authors read and approved the final manuscript.

