## Supplementary data including Figures S1, S2, S3, S4, S5, S6, and S7, Tables S1 and S2, and Sections S1, S2, and S3 are available. for "Umibato: estimation of time-varying microbial interaction using continuous-time regression hidden Markov model"

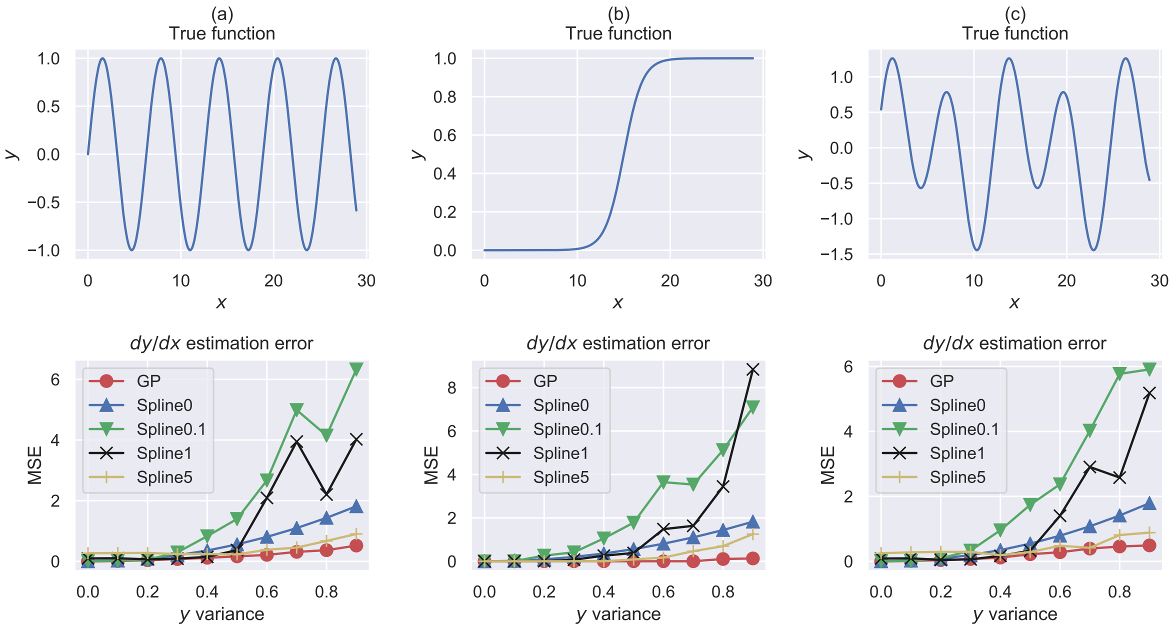


Figure S1: True function and the mean squared error (MSE) of growth rate estimation methods (Gaussian process regression, penalized spline) for synthetic data. The top row represents the shape of the true function, and the bottom row shows the MSE corresponding to the true function. The $x$- and $y$-axes in the top row represent $x$ and $y$, respectively. The lower $x$- and $y$-axes represent noise and MSE, respectively. "GP" represents the Gaussian process, and "Spline$p$" represents the penalized spline, where $p$ is the coefficient of the penalty term. Each column is the result of (a) $y = \sin(x)$, (b) $y = \text{sigmoid}(x - 15)$, and (c) $y = 0.5\sin(0.5x) + \cos(x - 1)$ being a true function.

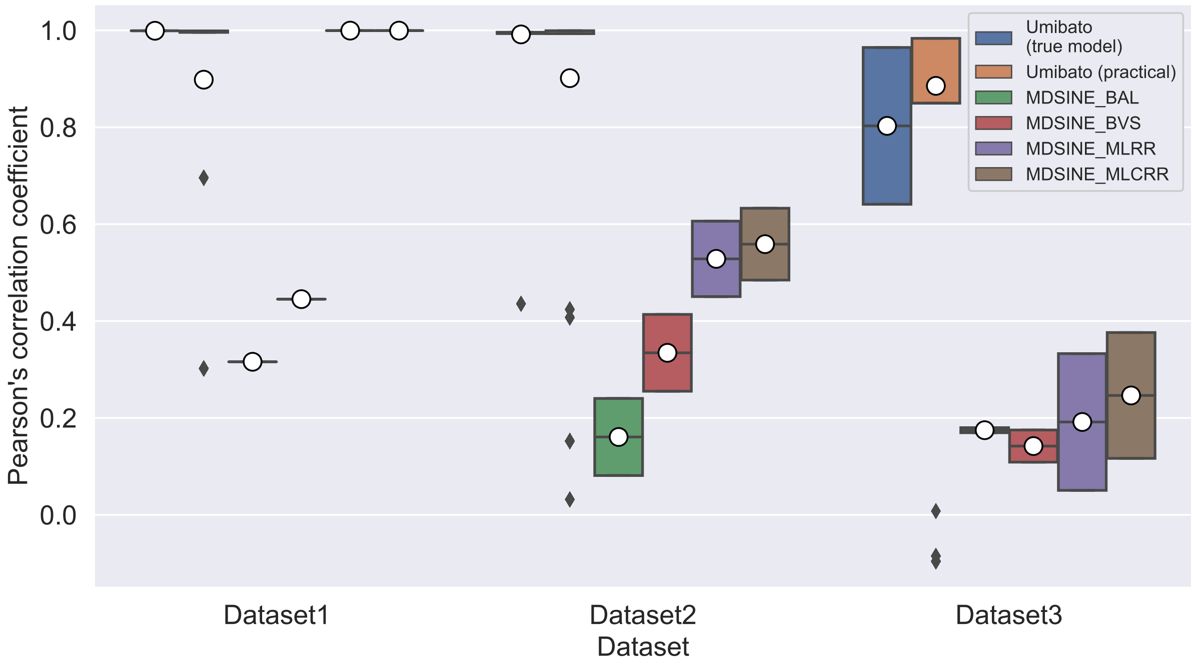


Figure S2: Box plot of Pearson's correlation coefficients between true parameters and parameters estimated by each method for each synthetic dataset. The $x$-axis indicates datasets, and the $y$-axis indicates the Pearson's correlation coefficients of the gLVE parameters of each observation point. The six boxes indicate Umibato in the true model case, Umibato in the practical case, BAL, BVS, MLRR, and MLCRR in order from left to right. Each white circle indicates the mean of the observation points for each method. Black points indicate outliers.

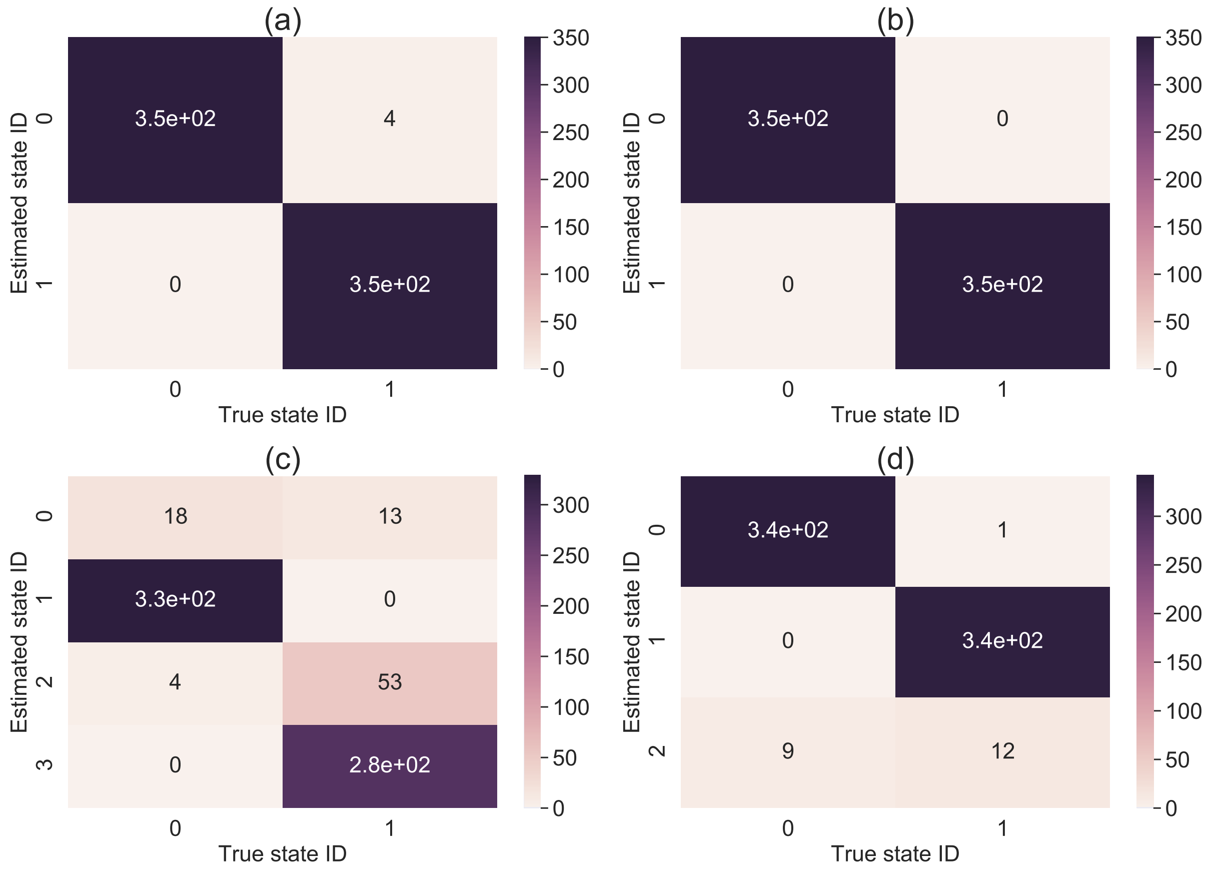


Figure S3: The cross-tabulation table between true interaction states and interaction states estimated by Umibato in each case on the multi-state synthetic datasets. The $x$- and $y$-axes represent the true and estimated state IDs, respectively. Darker colors indicate higher probabilities. **(a)** In the true model case of Dataset 2. **(b)**In the true model case of Dataset 3. **(c)** In the practical case of Dataset 2. **(d)**In the practical case of Dataset 3.

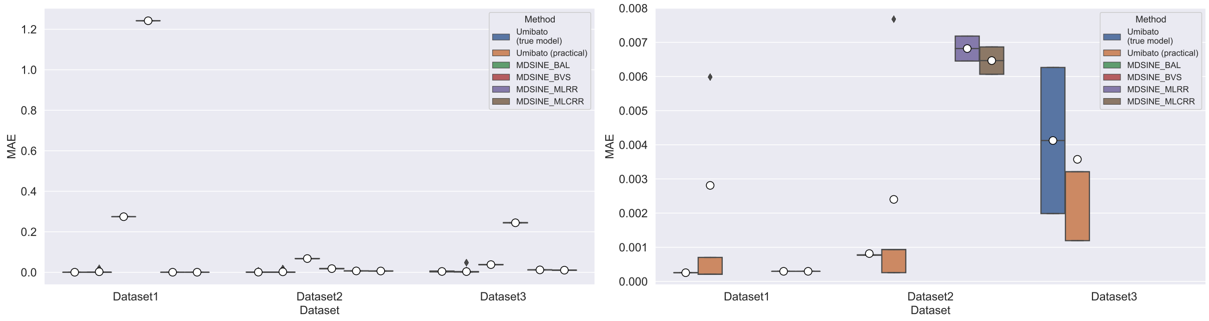


Figure S4: Mean absolute error (MAE) of gLVE parameters by each method estimation for each synthetic dataset. The $x$-axis indicates the datasets and the $y$-axis indicates the MAE. The six boxes indicate Umibato in the true model case, Umibato in the practical case, BAL, BVS, MLRR, and MLCRR in order from left to right. (**a**)Figure of the range containing all the values. (**b**) Figure of the range focused on small values.

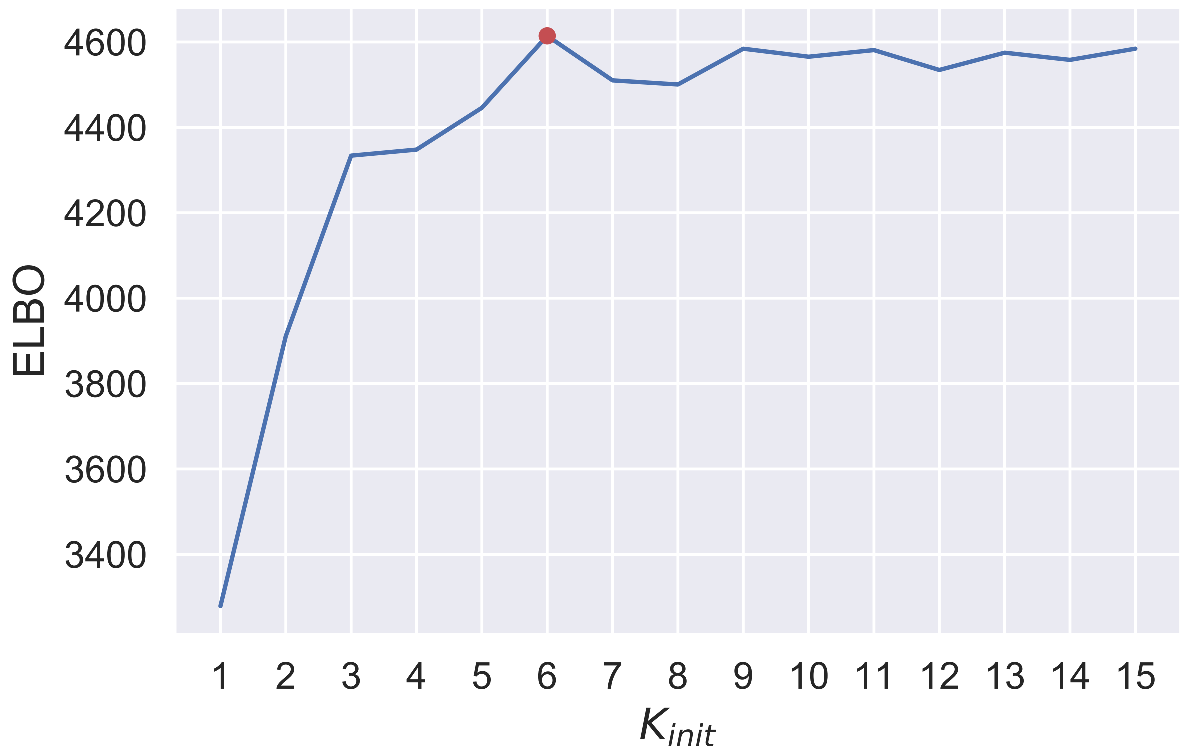


Figure S5: Maximized ELBO for each $K_{init}$. The $x$- and $y$-axes indicate $K_{init}$ and the maximum value of the maximized ELBO, respectively. A red point indicates the maximum value.

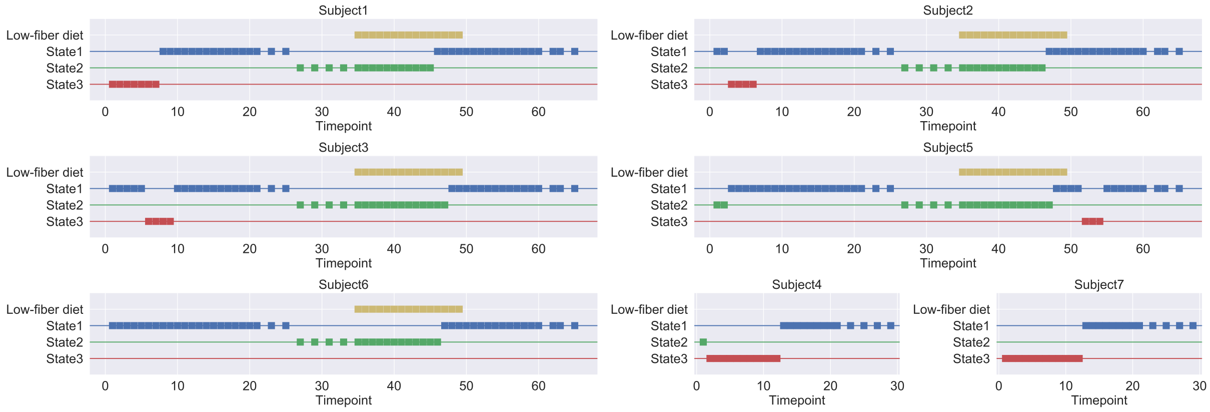


Figure S6: Estimated maximum likelihood path for each subject in the case $K_{init} = 3$.

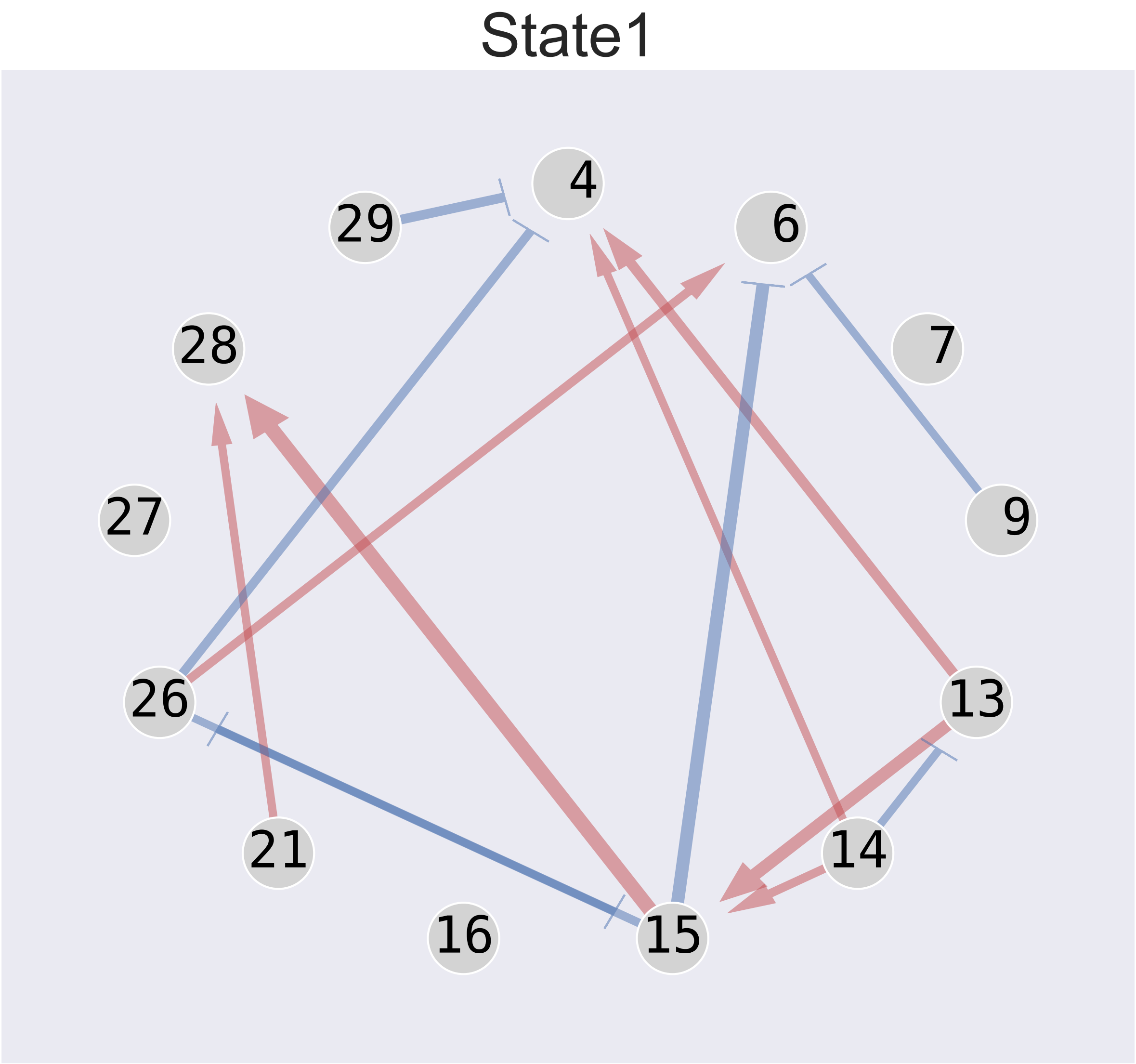


Figure S7: Bacterial network estimated by the single-state model.

Table S1: Variable notations.

| Variables | Representation/Definition |
|---|---|
| $\mathbf{X}$ | $\left(\mathbf{X}^{(1)}, \ldots, \mathbf{X}^{(s)}, \ldots, \mathbf{X}^{(S)}\right)^{\mathrm{T}}$ |
| $\mathbf{X}^{(s)}$ | $\left(\mathbf{x}_1^{(s)}, ..., \mathbf{x}_n^{(s)}, ..., \mathbf{x}_{N_s}^{(s)}\right)$ |
| $\mathbf{x}_n^{(s)}$ | $\left(1, x_{n,1}^{(s)}, ..., x_{n,i}^{(s)}, ..., x_{n,M}^{(s)}\right)^{\mathrm{T}}$ |
| $x_{n,i}^{(s)}$ | the quantitative abundance of $i$-th microbe of the $n$-th observation point of the $s$-th subject |
| $x_{n,0}^{(s)}$ | 1 (the element for the growth parameter) |
| $\mathbf{Y}$ | $\left(\mathbf{Y}^{(1)}, \ldots, \mathbf{Y}^{(s)}, \ldots, \mathbf{Y}^{(S)}\right)^{\mathrm{T}}$ |
| $\mathbf{Y}^{(s)}$ | $\left(\mathbf{y}_1^{(s)}, ..., \mathbf{y}_n^{(s)}, ..., \mathbf{y}_{N_s}^{(s)}\right)$ |
| $\mathbf{y}_n^{(s)}$ | $\left(y_{n,1}^{(s)}, ..., y_{n,i}^{(s)}, ..., y_{n,M}^{(s)}\right)^{\mathrm{T}}$ |
| $y_{n,i}^{(s)}$ | the growth rate of $i$-th microbe of the $n$-th observation point of the $s$-th subject |
| $\boldsymbol{\Sigma}$ | $\left(\boldsymbol{\Sigma}^{(1)}, \ldots, \boldsymbol{\Sigma}^{(s)}, \ldots, \boldsymbol{\Sigma}^{(S)}\right)^{\mathrm{T}}$ |
| $\boldsymbol{\Sigma}^{(s)}$ | $\left(\boldsymbol{\sigma}_1^{(s)}, ..., \boldsymbol{\sigma}_n^{(s)}, ..., \boldsymbol{\sigma}_{N_s}^{(s)}\right),$ |
| $\boldsymbol{\sigma}_n^{(s)}$ | $\left(\sigma_{n,1}^{(s)}, ..., \sigma_{n,i}^{(s)}, ..., \sigma_{n,M}^{(s)}\right)^{\mathrm{T}}$ |
| $\sigma_{n,i}^{(s)}$ | the variance of the growth rate of $i$-th microbe of the $n$-th observation point of the $s$-th subject |
| $\mathbf{Z}$ | $\left(\mathbf{Z}^{(1)}, \ldots, \mathbf{Z}^{(s)}, \ldots, \mathbf{Z}^{(S)}\right)^{\mathrm{T}}$ |
| $\mathbf{Z}^{(s)}$ | $\left(\mathbf{z}_1^{(s)}, ..., \mathbf{z}_n^{(s)}, ..., \mathbf{z}_{N_s}^{(s)}\right)$ |
| $\mathbf{z}_n^{(s)}$ | the one-hot vector that indicates the state of the $n$-th observation point of the $s$-th subject |
| $\boldsymbol{\zeta}(t)$ | $\left\{\boldsymbol{\zeta}^{(s)}(t)\right\}_{s=1}^{S}$ |
| $\boldsymbol{\zeta}^{(s)}(t)$ | $\left\{\boldsymbol{\zeta}_n^{(s)}(t)\right\}_{s=1}^{S}$ |
| $\boldsymbol{\zeta}_n^{(s)}(t)$ | the stochastic process of states between the $n$-th and $(n+1)$-th observation points of the $s$-th subject at time $t$ |
| $\boldsymbol{\Phi}$ | $\{\boldsymbol{\Phi}_k\}_{k=1}^{K}$ |
| $\boldsymbol{\Phi}_k$ | $\left(\boldsymbol{\phi}_{k,1}, ..., \boldsymbol{\phi}_{k,i}, ..., \boldsymbol{\phi}_{k,M}\right)^{\mathrm{T}}$ |
| $\boldsymbol{\phi}_{k,i}$ | $\left(\phi_{k,i,0}, ..., \phi_{k,i,j}, ..., \phi_{k,i,M}\right)^{\mathrm{T}}$ |
| $\phi_{k,i,0}$ | the growth parameter of the $i$-th microbe |
| $\phi_{k,i,j}$ | the interaction parameter from the $j$-th microbe to the $i$-th microbe |
| $\mathbf{d}$ | $\left\{\mathbf{d}^{(s)}\right\}_{s=1}^{S}$ |
| $\mathbf{d}^{(s)}$ | $\left\{d_n^{(s)}\right\}_{n=1}^{N_s-1}$ |
| $d_n^{(s)}$ | the time interval between $n$-th and $(n+1)$-th observation points of $s$-th subject |
| $\mathbf{Q}$ | the transition rate matrix |
| $q_{k,l}$ | the $(k,l)$ element of $\mathbf{Q}$ |
| $\mathbf{P}(t)$ | $\left(\mathbf{p}_1(t), ..., \mathbf{p}_k(t), ..., \mathbf{p}_K(t)\right)^{\mathrm{T}},$ |
| $\mathbf{p}_k(t)$ | $\left(p_{k,1}(t), ..., p_{k,k'}(t), ..., p_{k,K}(t)\right)^{\mathrm{T}},$ |
| $p_{k,l}(t)$ | the transition probability from the $k$-th state to the $l$-th state when $t$ units of time has elapsed |
| $\boldsymbol{\lambda}$ | $\{\lambda_i\}_{i=1}^{M}$ |
| $\lambda_i$ | Parameter of the prior distribution of $\phi_{k,i,j}$ |

Table S2: Lineages of the strains based on the study by Atarashi *et al.* (2013).

| Strain ID | Bacteria |
|---|---|
| 4 | *Clostridium hathewayi* |
| 6 | *Blautia producta* |
| 7 | *Clostridium bolteae* |
| 9 | *Clostridium indolis* |
| 13 | *Anaerotruncus colihominis* |
| 14 | *Ruminococcus sp. ID8* |
| 15 | *Clostridium asparagiforme* |
| 16 | *Clostridium 7 3 54FAA* |
| 21 | *Eubacterium fissicatena* |
| 26 | *Clostridium scindens* |
| 27 | *Lachnospiraceae 3_ 1_ 57FAA* |
| 28 | *Clostridiales 1_ 7_ 47FAA* |
| 29 | *Lachnospiraceae 3_ 1_ 57FAA* |

### S1 Derivation of variational inference algorithm

The notations of the variables are described in Table S1.

#### S1.1 Decompostion of ELBO

Decompose the ELBO according to the generative model as follows:

$$
\begin{aligned}
\mathcal{L} = &\Big\langle \sum_s \ln p(\mathbf{Y}^{(s)}, \boldsymbol{\zeta}^{(s)}(t), \mathbf{Z}^{(s)}, \boldsymbol{\Phi} | \mathbf{X}^{(s)}, \boldsymbol{\Sigma}^{(s)}, \eta, \mathbf{Q}, \mathbf{d}^{(s)}, \boldsymbol{\lambda}) \\
&- \ln q(\boldsymbol{\zeta}^{(s)}(t), \mathbf{Z}^{(s)}) \Big\rangle_{\zeta, \mathbf{Z}, \boldsymbol{\Phi}} - \langle \ln q(\boldsymbol{\Phi}) \rangle_{\boldsymbol{\Phi}} \\
&= \sum_s \Big\langle \ln p(\mathbf{Y}^{(s)} | \boldsymbol{\zeta}^{(s)}(t), \mathbf{Z}^{(s)}, \boldsymbol{\Phi}, \mathbf{X}^{(s)}, \boldsymbol{\Sigma}^{(s)}, \eta) \Big\rangle_{\mathbf{Z}, \boldsymbol{\Phi}} \\
&+ \sum_s \Big\langle \ln p(\boldsymbol{\zeta}^{(s)}(t) | \mathbf{Z}^{(s)}, \mathbf{Q}) \Big\rangle_{\zeta, \mathbf{Z}} \\
&+ \sum_s \Big\langle \ln p(\mathbf{Z}^{(s)} | \mathbf{Q}, \mathbf{d}^{(s)}) \Big\rangle_{\mathbf{Z}} \\
&+ \langle \ln p(\boldsymbol{\Phi} | \boldsymbol{\lambda}) \rangle_{\boldsymbol{\Phi}} - \langle \ln q(\zeta(t), \mathbf{Z}) \rangle_{\zeta, \mathbf{Z}} - \langle \ln q(\boldsymbol{\Phi}) \rangle_{\boldsymbol{\Phi}}.
\end{aligned}
$$

#### S1.2 Update of $q(\mathbf{\Phi})$

We obtained the partial derivative of the ELBO with respect to $q(\mathbf{\Phi})$ using the variational method as follows:

$$\frac{\partial \mathcal{L}}{\partial q(\mathbf{\Phi})} = \left\langle \ln p(\mathbf{Y}^{(s)}|\mathbf{Z}^{(s)}, \mathbf{\Phi}, \mathbf{X}^{(s)}, \mathbf{\Sigma}^{(s)}, \eta) \right\rangle_{\mathbf{Z}} + \ln p(\mathbf{\Phi}|\boldsymbol{\lambda}) - \ln q(\mathbf{\Phi}) - 1 = 0.$$

The following stationary point of the ELBO is then obtained:

$$\begin{aligned}
\ln q(\mathbf{\Phi}) &= \left\langle \ln p(\mathbf{Y}^{(s)}|\mathbf{Z}^{(s)}, \mathbf{\Phi}, \mathbf{X}^{(s)}, \mathbf{\Sigma}^{(s)}, \eta) \right\rangle_{\mathbf{Z}} + \ln p(\mathbf{\Phi}|\boldsymbol{\lambda}) + \text{const.} \\
&= \sum_s \sum_n \sum_k \left\langle z_{n,k}^{(s)} \right\rangle_{\mathbf{Z}} \ln p(\mathbf{y}_n^{(s)}|z_{n,k}^{(s)} = 1, \mathbf{\Phi}_k, \mathbf{x}_n^{(s)}, \mathbf{\Sigma}^{(s)}, \eta) + \sum_k \sum_i \ln p(\boldsymbol{\phi}_{k,i}|\lambda_m) + \text{const.}
\end{aligned}$$

We can decompose $\ln q(\mathbf{\Phi})$ into $\ln q(\boldsymbol{\phi}_{k,i})$ as follows:

$$\begin{aligned}
\ln q(\boldsymbol{\phi}_{k,i}) &= \sum_s \sum_n \left\langle z_{n,k}^{(s)} \right\rangle_{z_{n,k}^{(s)}} \ln p(y_{n,i}^{(s)}|z_{n,k}^{(s)} = 1, \boldsymbol{\phi}_{k,i}, x_{n,i}^{(s)}, \sigma_{n,i}^{(s)}, \eta) + \ln p(\boldsymbol{\phi}_{k,i}|\lambda_m) + \text{const.} \\
&= \sum_s \sum_n \gamma_{n,k}^{(s)} \ln \text{Normal}\Big(y_{n,i}^{(s)}|{\boldsymbol{\phi}_{k,i}}^{\mathrm{T}} \mathbf{x}_n^{(s)}, \eta\sigma_{n,i}^{(s)}\Big) + \ln \text{MultiNormal}\big(\boldsymbol{\phi}_{k,i}|\mathbf{0}, \lambda_m \mathbf{I}\big) + \text{const.}
\end{aligned}$$

Here,

$$\begin{aligned}
\sum_s \sum_n \gamma_{n,k} \ln \text{Normal}\Big(y_{n,i}^{(s)}|{\mathbf{x}_n^{(s)}}^{\mathrm{T}} \boldsymbol{\phi}_{k,i}, \eta\sigma_{n,i}^{(s)}\Big) &= \frac{1}{2}\Big(\mathbf{y}_{\cdot,i}^{(\cdot)} - \mathbf{X}\boldsymbol{\phi}_{k,i}\Big)^{\mathrm{T}} \mathbf{L}_{k,i} \Big(\mathbf{y}_{\cdot,i}^{(\cdot)} - \mathbf{X}\boldsymbol{\phi}_{k,i}\Big) + \text{const.} \\
&= \ln \text{MultiNormal}\Big(\mathbf{y}_{\cdot,i}^{(\cdot)}|\mathbf{X}\boldsymbol{\phi}_{k,i}, \mathbf{L}_{k,i}\Big),
\end{aligned}$$

where

$$\mathbf{L}_{k,i} = \text{diag}(\boldsymbol{\gamma}_{\cdot,k}^{(\cdot)})\text{diag}(\eta\boldsymbol{\sigma}_{\cdot,i}^{(\cdot)})^{-1}.$$

Then, we obtained the following:

$$\ln q(\boldsymbol{\phi}_{k,i}) = \ln \text{MultiNormal}\Big(\mathbf{y}_{\cdot,i}^{(\cdot)}|\mathbf{X}\boldsymbol{\phi}_{k,i}, \mathbf{L}_{k,i}\Big) + \ln \text{MultiNormal}\big(\boldsymbol{\phi}_{k,i}|\mathbf{0}, \lambda_i \mathbf{I}\big) + \text{const.}$$

Finally, $q(\boldsymbol{\phi}_{k,i})$ can be represented the following distribution by normalization:

$$q(\boldsymbol{\phi}_{k,i}) = \text{MultiNormal}\Big(\boldsymbol{\phi}_{k,i}|\mathbf{\Sigma}_{\boldsymbol{\phi}_{k,i}} \mathbf{t}_{\boldsymbol{\phi}_{k,i}}, \mathbf{\Sigma}_{\boldsymbol{\phi}_{k,i}}\Big),$$

where

$$\begin{aligned}
\mathbf{t}_{\boldsymbol{\phi}_{k,i}} &= \mathbf{X}^{\mathrm{T}} \mathbf{L}_{k,i} \mathbf{y}_{\cdot,i}^{(\cdot)}, \\
\mathbf{\Sigma}_{\boldsymbol{\phi}_{k,i}} &= \Big(\lambda_i \mathbf{I} + \mathbf{X}^{\mathrm{T}} \mathbf{L}_{k,i} \mathbf{X}\Big)^{-1}.
\end{aligned}$$

##### S1.3 Update of $q(\mathbf{Z}^{(s)})$

In the same way as $q(\mathbf{\Phi})$, we obtained the following formula using the variational method:

$$\begin{aligned}\ln q(\mathbf{Z}^{(s)}) &= \left\langle \ln p(\mathbf{Y}^{(s)}|\mathbf{Z}^{(s)}, \mathbf{\Phi}, \mathbf{X}^{(s)}, \mathbf{\Sigma}^{(s)}, \eta) \right\rangle_{\mathbf{\Phi}} + \ln p(\mathbf{Z}^{(s)}|\mathbf{Q}, \mathbf{d}^{(s)}) - \ln \tilde{p}(\mathbf{Y}^{(s)}) \\ &= \sum_n \sum_i \left\langle \ln p(y_{n,i}^{(s)}|z_{n,k}^{(s)} = 1, \boldsymbol{\phi}_{k,i}, \mathbf{x}_n^{(s)}, \sigma_{n,i}^{(s)}, \eta) \right\rangle_{\boldsymbol{\phi}_{k,i}} + \sum_n \ln p(\mathbf{z}_{n+1}^{(s)}|\mathbf{z}_n^{(s)}, \mathbf{Q}, d_n^{(s)}) - \ln \tilde{p}(\mathbf{Y}^{(s)}),\end{aligned}$$

where

$$\ln \tilde{p}(\mathbf{Y}^{(s)}) = \ln \sum_{\mathbf{Z}} \exp\left( \left\langle \ln p(\mathbf{Y}^{(s)}|\mathbf{Z}^{(s)}, \mathbf{\Phi}, \mathbf{X}^{(s)}, \mathbf{\Sigma}^{(s)}, \eta) \right\rangle_{\mathbf{\Phi}} + \ln p(\mathbf{Z}^{(s)}|\mathbf{Q}, \mathbf{d}^{(s)}) \right),$$

$$\sum_n \ln p(\mathbf{z}_{n+1}^{(s)}|\mathbf{z}_n^{(s)}, \mathbf{Q}, d_n^{(s)}) = -\sum_k z_{1,k}^{(s)} \ln K + \sum_n^{N_s-1} \sum_{k,l} z_{n,k}^{(s)} z_{n+1,l}^{(s)} \ln\left(p_{k,l}(d_n^{(s)})\right). \tag{S1}$$

The $q(\boldsymbol{\phi}_{k,i})$ expectation term can be written as

$$\begin{aligned}&\left\langle \ln p(y_{n,i}^{(s)}|z_{n,k}^{(s)} = 1, \boldsymbol{\phi}_{k,i}, \mathbf{x}_n^{(s)}, \sigma_{n,i}^{(s)}, \eta) \right\rangle_{\boldsymbol{\phi}_{k,i}} \\ &= -\frac{1}{2} \ln 2\pi\eta {\sigma_{n,i}^{(s)}}^2 - \left\langle \frac{1}{2\eta {\sigma_{n,i}^{(s)}}^2} \left\{ {y_{n,i}^{(s)}}^2 - 2y_{n,i}^{(s)} {\mathbf{x}_n^{(s)}}^{\mathrm{T}} \boldsymbol{\phi}_{k,i} + {\mathbf{x}_n^{(s)}}^{\mathrm{T}} \boldsymbol{\phi}_{k,i} {\boldsymbol{\phi}_{k,i}}^{\mathrm{T}} \mathbf{x}_n^{(s)}) \right\} \right\rangle_{\boldsymbol{\phi}_{k,i}} \\ &= -\frac{1}{2} \ln 2\pi\eta {\sigma_{n,i}^{(s)}}^2 - \frac{1}{2\eta {\sigma_{n,i}^{(s)}}^2} \left\{ {y_{n,i}^{(s)}}^2 - 2y_{n,i}^{(s)} {\mathbf{x}_n^{(s)}}^{\mathrm{T}} \langle \boldsymbol{\phi}_{k,i} \rangle_{\boldsymbol{\phi}_{k,i}} + {\mathbf{x}_n^{(s)}}^{\mathrm{T}} \left\langle \boldsymbol{\phi}_{k,i} {\boldsymbol{\phi}_{k,i}}^{\mathrm{T}} \right\rangle_{\boldsymbol{\phi}_{k,i}} \mathbf{x}_n^{(s)}) \right\}.\end{aligned}$$

#### S1.4 Update of $q(\boldsymbol{\zeta}(t))$

We used the true $p(\boldsymbol{\zeta}^{(s)}(t)|\mathbf{Z}^{(s)}, \mathbf{Q}, \mathbf{d})$ as $q(\boldsymbol{\zeta}^{(s)}(t)|\mathbf{Z}^{(s)})$. $q(\boldsymbol{\zeta}^{(s)}(t)|\mathbf{Z}^{(s)})$ can be decomposed as follows:

$$
\begin{aligned}
q(\boldsymbol{\zeta}^{(s)}(t)|\mathbf{Z}^{(s)}) &= p(\boldsymbol{\zeta}^{(s)}(t)|\mathbf{Z}^{(s)}, \mathbf{Q}, \mathbf{d}) \\
&= \prod_{n=1}^{N_s-1} \prod_{u=1}^{K} \prod_{v=1}^{K} p(\boldsymbol{\zeta}_n^{(s)}(t)|z_{n,u}^{(s)} = 1, z_{n+1,v}^{(s)} = 1, \mathbf{Q}, d_n^{(s)})^{z_{n,u}^{(s)} z_{n+1,v}^{(s)}} \\
&= \prod_{n=1}^{N_s-1} \prod_{u=1}^{K} \prod_{v=1}^{K} \prod_{k=1}^{K} c(n,s,t,k,u,v)^{\zeta_{n,k}^{(s)}(t) z_{n,u}^{(s)} z_{n+1,v}^{(s)}}.
\end{aligned} \tag{S2}
$$

Here, $c(n,s,t,k,u,v)$ is represented as follows:

$$
\begin{aligned}
c(n,s,t,k,u,v) &= p(\zeta_{n,k}^{(s)}(t) = 1|z_{n,u}^{(s)} = 1, z_{n+1,v}^{(s)} = 1, \mathbf{Q}, d_n^{(s)}) \\
&= \frac{p(\zeta_{n,k}^{(s)}(t) = 1, z_{n+1,v}^{(s)} = 1|z_{n,u}^{(s)} = 1, \mathbf{Q}, d_n^{(s)})}{p(z_{n+1,v}^{(s)} = 1|z_{n,u}^{(s)} = 1, \mathbf{Q}, d_n^{(s)})} \\
&= \frac{p_{u,k}(t) p_{k,v}(d_n^{(s)} - t)}{p_{u,v}(d_n^{(s)})}.
\end{aligned}
$$

Therefore, we can represent the expectation values of $\tau_k^{(s,n,u,v)}$ and $\nu_{k,k'}^{(s,n,u,v)}$ as follows:

$$
\begin{aligned}
\left\langle \tau_k^{(s,n,u,v)} \right\rangle_{\boldsymbol{\zeta}_n^{(s)}(t)} &= \frac{1}{p_{u,v}(d_n^{(s)})} \int_0^{d_n^{(s)}} p_{u,k}(t) p_{k,v}(d_n^{(s)} - t) dt, \\
\left\langle \nu_{k,k'}^{(s,n,u,v)} \right\rangle_{\boldsymbol{\zeta}_n^{(s)}(t)} &= \frac{q_{k,k'}}{p_{u,v}(d_n^{(s)})} \int_0^{d_n^{(s)}} p_{u,k}(t) p_{k',v}(d_n^{(s)} - t) dt.
\end{aligned}
$$

The calculation of these integrals was based on the study of Liu *et al.* (2015) and is given as follows:

$$
\int_0^{d_n^{(s)}} p_{u,k}(t) p_{k',v}(d_n^{(s)} - t) dt = \mathbf{exp}(\mathbf{A}t)_{1:K,K+1:2K},
$$

where

$$
\begin{aligned}
\mathbf{A} &= \begin{pmatrix} \mathbf{Q} & \mathbf{B} \\ \mathbf{0} & \mathbf{Q} \end{pmatrix}, \\
\mathbf{B} &= \mathbf{I}(k,k').
\end{aligned}
$$

Here, $\mathbf{0}$ is a matrix with 0 in the all elements, and $\mathbf{I}(i,j)$ is a matrix with 1 in the $(i,j)$ element and 0 otherwise.

##### S1.5 Update of maximum likelihood estimated parameters

We derived the updates for the maximum likelihood estimated parameters $\mathbf{Q}$, $\boldsymbol{\lambda}$, and $\eta$. The prior distribution for $\boldsymbol{\zeta}(t)$ and $\mathbf{Z}$ can be rewritten by

$$\ln p(\boldsymbol{\zeta}(t), \mathbf{Z}|\mathbf{Q}, \mathbf{d}) = \sum_k \left( \sum_{k' \neq k} \nu_{k,k'} \ln q_{k,k'} \right) - \left( \sum_{k'} q_{k,k'} \right) \tau_k.$$

As such, we obtained the update of $\mathbf{Q}$ as follows:

$$\begin{aligned} \frac{\partial \mathcal{L}}{\partial q_{k,k'}} &= \frac{\langle \nu_{k,k'} \rangle_{\boldsymbol{\zeta}(t),\mathbf{Z}}}{q_{k,k'}} - \langle \tau_k \rangle_{\boldsymbol{\zeta}(t),\mathbf{Z}} = 0, \\ q_{k,k'} &= \frac{\langle \nu_{k,k'} \rangle_{\boldsymbol{\zeta}(t),\mathbf{Z}}}{\langle \tau_k \rangle_{\boldsymbol{\zeta}(t),\mathbf{Z}}}. \end{aligned}$$

We obtained the update of $\boldsymbol{\lambda}$ as follows:

$$\begin{aligned} \frac{\partial \mathcal{L}}{\partial \lambda_m} &= \sum_k \frac{\partial}{\partial \lambda_m} \langle \ln p(\boldsymbol{\phi}_{k,i}|\lambda_m) \rangle_{\boldsymbol{\phi}_{k,i}} \\ &= \sum_k \frac{M+1}{2\lambda_m} - \frac{1}{2} \left\langle {\boldsymbol{\phi}_{k,i}}^{\mathrm{T}} \boldsymbol{\phi}_{k,i} \right\rangle_{\boldsymbol{\phi}_{k,i}} = 0, \\ \frac{(M+1)K}{2\lambda_m} &= \frac{1}{2} \sum_k \left\langle {\boldsymbol{\phi}_{k,i}}^{\mathrm{T}} \boldsymbol{\phi}_{k,i} \right\rangle_{\boldsymbol{\phi}_{k,i}}, \\ \lambda_m &= \frac{(M+1)K}{\sum_k \left\langle {\boldsymbol{\phi}_{k,i}}^{\mathrm{T}} \boldsymbol{\phi}_{k,i} \right\rangle_{\boldsymbol{\phi}_{k,i}}}. \end{aligned}$$

We obtained the update of $\eta$ as follows:

$$\begin{aligned} \frac{\partial \mathcal{L}}{\partial \eta} &= \frac{\partial}{\partial \eta} \left\langle \ln p(\mathbf{Y}^{(s)}|\mathbf{Z}^{(s)}, \boldsymbol{\Phi}, \mathbf{X}^{(s)}, \boldsymbol{\Sigma}^{(s)}, \eta) \right\rangle_{\mathbf{Z},\boldsymbol{\Phi}} \\ &= \sum_s \sum_n \sum_i \sum_k \frac{\partial}{\partial \eta} \left\{ -\frac{\gamma_{n,k}}{2} \ln \eta {\sigma_{n,i}^{(s)}}^2 - \frac{\gamma_{n,k}}{2\eta {\sigma_{n,i}^{(s)}}^2} \left\langle \left( y_{n,m} - {\mathbf{x}_n^{(s)}}^{\mathrm{T}} \boldsymbol{\phi}_{k,i} \right)^2 \right\rangle_{\boldsymbol{\phi}_{k,i}} \right\} \\ &= \sum_s \sum_n \sum_i \sum_k \left\{ -\frac{\gamma_{n,k}}{2\eta} + \frac{\gamma_{n,k}}{2\eta^2 \sigma_{n,i}^{(s)}} \left\langle \left( y_{n,m} - {\mathbf{x}_n^{(s)}}^{\mathrm{T}} \boldsymbol{\phi}_{k,i} \right)^2 \right\rangle_{\boldsymbol{\phi}_{k,i}} \right\} \\ &= -\frac{NM}{2\eta} + \frac{1}{2\eta^2} \sum_s \sum_n \sum_i \sum_k \frac{\gamma_{n,k}}{\sigma_{n,i}^{(s)}} \left\langle \left( y_{n,m} - {\mathbf{x}_n^{(s)}}^{\mathrm{T}} \boldsymbol{\phi}_{k,i} \right)^2 \right\rangle_{\boldsymbol{\phi}_{k,i}} = 0, \\ \eta &= \frac{1}{NM} \sum_s \sum_n \sum_i \sum_k \frac{\gamma_{n,k}}{\sigma_{n,i}^{(s)}} \left\langle \left( y_{n,m} - {\mathbf{x}_n^{(s)}}^{\mathrm{T}} \boldsymbol{\phi}_{k,i} \right)^2 \right\rangle_{\boldsymbol{\phi}_{k,i}}. \end{aligned}$$

##### S1.6 Calculation of ELBO

ELBO is obtained as follows:

$$\mathcal{L} = \left( \sum_s \left\langle \ln p(\mathbf{Y}^{(s)}, \boldsymbol{\zeta}^{(s)}(t), \mathbf{Z}^{(s)}, |\mathbf{\Phi}, \mathbf{X}^{(s)}, \mathbf{\Sigma}^{(s)}, \mathbf{Q}, \mathbf{d}, \eta) \right\rangle_{\boldsymbol{\zeta}^{(s)}(t), \mathbf{Z}^{(s)}, \mathbf{\Phi}} + \mathrm{H}[q(\boldsymbol{\zeta}^{(s)}(t), \mathbf{Z}^{(s)})] \right)$$
$$+ \left( \sum_k \sum_i \left\langle \ln p(\boldsymbol{\phi}_{k,i} | \lambda_m) \right\rangle_{\boldsymbol{\phi}_{k,i}} + \mathrm{H}[q(\boldsymbol{\phi}_{k,i})] \right),$$

where H[.] is the entropy of probabilistic distributions. The first term is transformed as follows:

$$\left\langle \ln p(\mathbf{Y}^{(s)}, \boldsymbol{\zeta}^{(s)}(t), \mathbf{Z}^{(s)} | \mathbf{\Phi}, \mathbf{X}^{(s)}, \mathbf{\Sigma}^{(s)}, \mathbf{Q}, \mathbf{d}, \eta) \right\rangle_{\boldsymbol{\zeta}^{(s)}(t), \mathbf{Z}^{(s)}, \mathbf{\Phi}} + \mathrm{H}[q(\boldsymbol{\zeta}^{(s)}(t), \mathbf{Z}^{(s)})]$$
$$= \left\langle \ln p(\mathbf{Y}^{(s)}, \mathbf{Z}^{(s)} | \mathbf{\Phi}, \mathbf{X}^{(s)}, \mathbf{\Sigma}^{(s)}, \mathbf{Q}, \mathbf{d}, \eta) \right\rangle_{\mathbf{Z}^{(s)}, \mathbf{\Phi}} + \left\langle \ln p(\boldsymbol{\zeta}^{(s)}(t) | \mathbf{Z}^{(s)}, \mathbf{Q}, \mathbf{d}) \right\rangle_{\boldsymbol{\zeta}^{(s)}(t), \mathbf{Z}^{(s)}}$$
$$+ \mathrm{H}[q(\mathbf{Z}^{(s)})] + \mathrm{H}[q(\boldsymbol{\zeta}^{(s)}(t) | \mathbf{Z}^{(s)})].$$

After updating $q(\mathbf{Z})$, we can substitute the following equation from Eq. (S1):

$$\left\langle \ln p(\mathbf{Y}^{(s)}, \mathbf{Z}^{(s)} | \mathbf{\Phi}, \mathbf{X}^{(s)}, \mathbf{\Sigma}^{(s)}, \mathbf{Q}, \mathbf{d}, \eta) \right\rangle_{\mathbf{Z}^{(s)}, \mathbf{\Phi}} + \mathrm{H}[q(\mathbf{Z}^{(s)})] = \left\langle \ln \tilde{p}(\mathbf{Y}^{(s)}) \right\rangle_{\mathbf{Z}^{(s)}}.$$

In addition, we can substitute the following equation from Eq. (S2):

$$\left\langle \ln p(\boldsymbol{\zeta}^{(s)}(t) | \mathbf{Z}^{(s)}, \mathbf{Q}, \mathbf{d}) \right\rangle_{\boldsymbol{\zeta}^{(s)}(t), \mathbf{Z}^{(s)}} + \mathrm{H}[q(\boldsymbol{\zeta}^{(s)}(t) | \mathbf{Z}^{(s)})] = 0.$$

We obtained

$$\left\langle \ln p(\mathbf{Y}^{(s)}, \boldsymbol{\zeta}^{(s)}(t), \mathbf{Z}^{(s)}, |\mathbf{\Phi}, \mathbf{X}^{(s)}, \mathbf{\Sigma}^{(s)}, \mathbf{Q}, \mathbf{d}, \eta) \right\rangle_{\boldsymbol{\zeta}^{(s)}(t), \mathbf{Z}^{(s)}, \mathbf{\Phi}} + \mathrm{H}[q(\boldsymbol{\zeta}^{(s)}(t), \mathbf{Z}^{(s)})]$$
$$= \left\langle \ln \tilde{p}(\mathbf{Y}^{(s)}) \right\rangle_{\mathbf{Z}^{(s)}} + 0 = \ln \tilde{p}(\mathbf{Y}^{(s)}).$$

Thus, ELBO is given by

$$\mathcal{L} = \sum_s \ln \tilde{p}(\mathbf{Y}^{(s)}) + \left( \sum_k \sum_i \left\langle \ln p(\boldsymbol{\phi}_{k,i} | \lambda_m) \right\rangle_{\boldsymbol{\phi}_{k,i}} + \frac{1}{2} \ln 2\pi e \Sigma_{\boldsymbol{\phi}_{k,i}} \right).$$

#### S2 Correction of gLVE parameters estimated using standardization of X

The parameters of gLVE can be obtained by correcting $\mathbf{\Phi}$ estimated from the standardized $\mathbf{X}$ using the following equation:

$$\hat{\phi}_{k,i,j} = \begin{cases} \frac{\tilde{\phi}_{k,i,j}}{\sigma_j^x} & (j > 0) \\ \tilde{\phi}_{k,i,j} - \sum_{j=1}^{M} \hat{\phi}_{k,i,j} \frac{\mu_j^x}{\sigma_j^x} & (j = 0), \end{cases}$$

where $\tilde{\phi}_{k,i,j}$ is the gLVE parameter corresponding to the $k$-th state estimated from standardized $\mathbf{X}$, $\hat{\phi}_{k,i,j}$ is the corrected $\tilde{\phi}_{k,i,j}$, and $\sigma_j^x$ and $\mu_j^x$ are the standard deviation and mean of the $j$-th microbe abundance, respectively.

#### S3 Synthetic data experiments

We created several datasets of 100 days based on gLVE. The gLVE parameters were randomly generated by the following generative process:

$$\phi_{k,i,j} \sim \begin{cases} \text{NegativeHalfNormal}(0, 0.01) & (i = j) \\ \text{PositiveHalfNormal}(0, 0.01) & (j = 0) \\ \text{Normal}(0, 0.01) & (otherwise) \end{cases}$$

The generative process of the gLVE parameters satisfies the assumptions in MDSINE (Bucci *et al.*, 2016). We randomly obtained the initial microbial quantitative abundance $\mathbf{x}(0)$ from Gamma$(1, 1)$ and computed $\mathbf{y}(0)$ using gLVE, where Gamma$(k, \theta)$ denotes the gamma distribution with shape parameter $k$ and scale parameter $\theta$. From this value, we iterated the gLVE and obtained the following equation:

$$\mathbf{x}(t + dt) = \exp(\ln \mathbf{x}(t) + \mathbf{y}(t)dt),$$

where $dt$ is a small time (0.001 days). The state switches from state 0 to state 1 in the middle of this period. $S = 7$, $d_n^{(s)} = 1$, and $N_s = 100$ were used in all datasets. We set different numbers of states $K$ and microbes $M$ for each dataset. Table S3 lists the settings for each dataset. We generated datasets until a non-divergent trajectory was obtained. Finally, we added random noises Normal$(0, 0.01)$ to generate $\mathbf{x}(t)$.

Table S3: Settings for each dataset.

| Dataset | $K$ | $M$ |
|---|---|---|
| Dataset1 | 1 | 5 |
| Dataset2 | 2 | 5 |
| Dataset3 | 2 | 10 |

#### S4 Computational time

We measured the computational time of the GPR and CTRHMM estimation on the mouse gut microbiome dataset. We measured the time of one GPR estimation and the average time of 10 CTRHMM trials. It took 7.86 and 24.3 seconds to fit the GPR and the CTRHMM, respectively. We used a ZenBook 3 UX390UA with a 2.7 GHz 2-core Intel Core i7 processor, 16GB of RAM, and Ubuntu 18.04 for measuring the computational time.
